# Dual recognition of phage SSB triggers DRT4 for antiphage defense

**DOI:** 10.64898/2026.10.21.760695

**Authors:** Monika Jasnauskaitė, Augustas Cibulskis, Kristina Kazlauskaitė, Aistė Skorupskaitė, Jonas Juozapaitis, Algirdas Mikšys, Miglė Tomkuvienė, Lina Malinauskaitė, Patrick Pausch

## Abstract

Defense-associated reverse transcriptase 4 (DRT4) protects bacteria against phages through single-stranded DNA (ssDNA) synthesis and activation of an effector RNase. How precisely phage recognition is coupled to RNase activation remains unclear. Here, we combine structural, biochemical, and genetic analyses to define the requirements for DRT4 activation. Cryo-electron microscopy structures capture DNA synthesis and provide a structural basis for preferential purine incorporation. Screening against the BASEL phage collection reveals protection against diverse phages and identifies the Bas17-encoded ssDNA-binding (SSB) protein gp44 as a DRT4 trigger. We show that DRT4 requires both recognition of its ssDNA product and direct protein-protein interactions of gp44 for RNase activation. Together, our findings support a previously unrecognized “two-factor authentication” model in which ssDNA-mediated recruitment of a phage SSB cooperates with specific protein interactions to license abortive infection for antiphage defense.

## Introduction

Bacteria have evolved diverse defense mechanisms to counter bacteriophage infection ^1^. Among these, abortive infection (Abi) systems induce growth arrest or programmed cell death in infected cells, limiting phage propagation and protecting the surrounding bacterial population ^2,3^. A growing class of Abi systems relies on reverse transcriptases (RTs), enzymes typically associated with RNA-templated DNA synthesis but increasingly acknowledged as versatile components of bacterial immunity ^4,5,6^.

Bacterial RTs are broadly classified into group II introns, retrons, diversity-generating retroelements (DGRs), and the abortive infection/unknown group (UG/Abi) ^7^. Based on gene organization and associated effector domains, UG/Abi systems are further divided into three major classes. Class 1 RTs are fused to C-terminal α-helical repeat (α-Rep) domains; class 2 RTs lack recognizable additional domains; and class 3 RTs are typically associated with or fused to C-N hydrolase or phosphohydrolase domains ^7^. Unlike canonical reverse transcriptases, UG/Abi class 1 enzymes, including AbiK, Abi-P2, AbiA, DRT4 (UG15) and DRT6 (UG12), catalyze template-independent DNA synthesis through a protein-priming mechanism, in which the first nucleotide is covalently attached to a tyrosine, threonine or serine residue ^8–12^. Recent cryo-EM structures of AbiK, DRT4 and DRT6 revealed conserved hexameric assemblies comprising two back-to-back trimers and captured covalently attached ssDNA products, providing insight into the mechanism of template-independent DNA synthesis ^8,10–12^.

The α-Rep domain is a defining feature of UG/Abi class 1 enzymes ^7^. These domains share a common architecture with a variable number of α-helical repeats despite lacking detectable sequence similarity. Structural studies show that the α-Rep domain occupies the position of the thumb domain in canonical RTs ^8,11,13^, yet its precise function remains unclear. Deleting just 46 C-terminal amino acids from AbiK abolishes antiphage defense ^8^, implicating the α-Rep domain in immune function. The close phylogenetic relationship between AbiK and DRT4 raises the possibility that the corresponding region also contributes to DRT4-mediated defense. However, whether the α-Rep domain serves structural purposes, mediates allosteric regulation, or senses phage-derived activators has not been experimentally established.

Recent studies have begun to uncover how UG/Abi systems are activated during phage infection ^14^. Phage-encoded single-stranded (ss)DNA-binding (SSB) proteins have emerged as a recurring class of immune response activators, and DRT4 has been shown to respond to the T5 phage SSB ORF55 ^11^. ORF55 was proposed to stabilize the DRT4-synthesized ssDNA, promoting DNA product accumulation and activation of the defense pathway ^11^. More recently, T7 SSB, together with elevated intracellular dNTP concentrations during phage infection, was proposed to activate DRT4’s guanosine-specific RNase activity. This activity cleaves both phage and host RNAs, ultimately inducing abortive infection ^12^. However, the molecular mechanism linking phage SSB recognition to DRT4 activation remains unresolved (**Fig. 1A**). In particular, whether the α-Rep domain contributes to phage trigger recognition or the regulation of RNase activity is unknown.

**Fig. 1.**
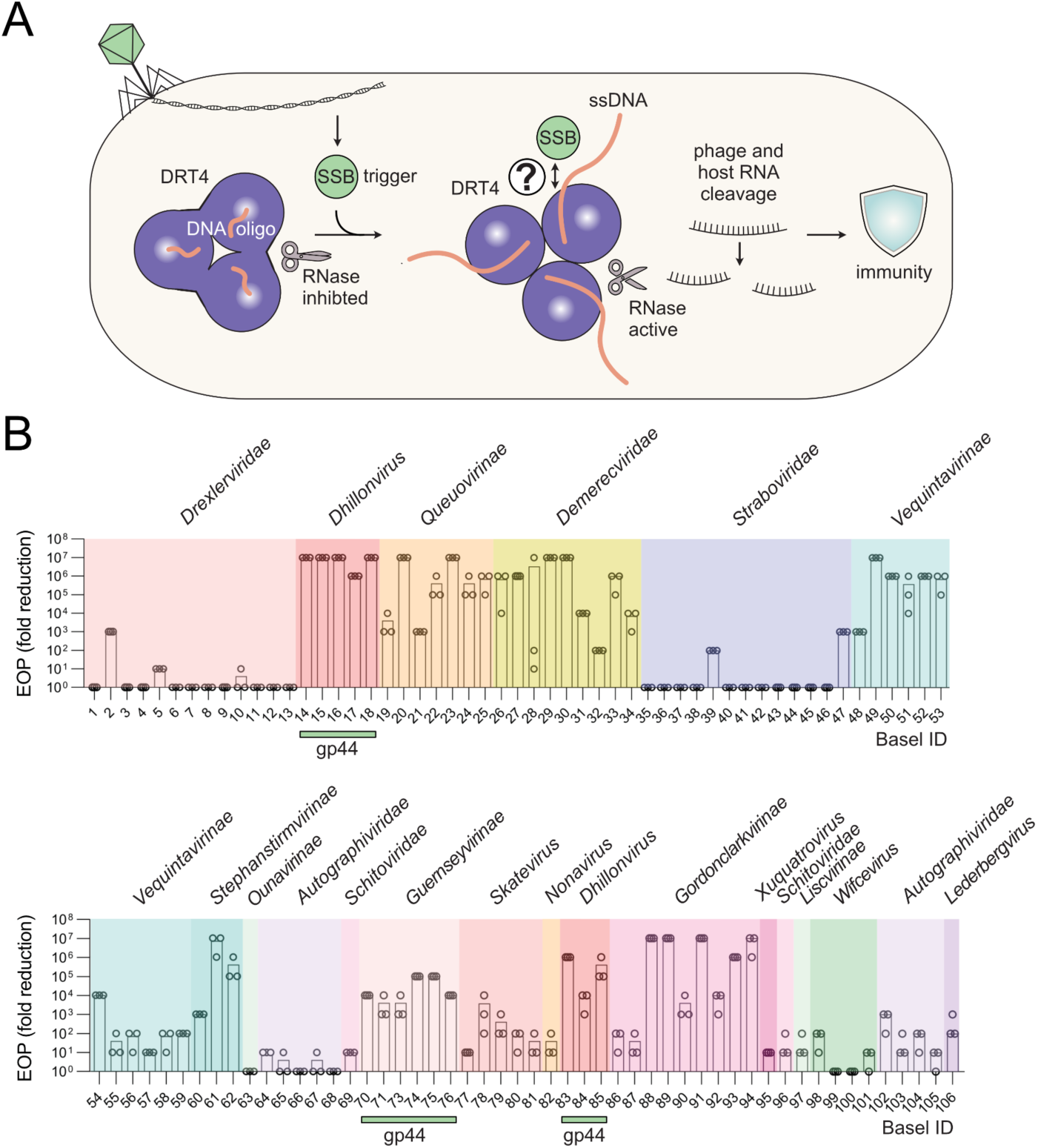
DRT4 protects against diverse bacteriophages. A, Schematic illustrating DRT4-mediated antiphage defense and the unresolved mechanism of activation by phage SSB proteins (question mark). **B,** Antiphage activity of DRT4 against BASEL phage collection, measured by efficiency of plating (EOP) compared to an empty plasmid control. *n* = 3 technical replicates; mean. Green bars indicate phages encoding gp44-homologs.

Here, we define the mechanism of SSB-dependent DRT4 activation. An expanded screen against the BASEL phage library revealed that DRT4 protects against diverse bacteriophages, and sequencing of escape phages identified the Bas17-encoded SSB protein gp44 as a DRT4 activator. Biochemical and genetic analyses showed that gp44 binds both DRT4-produced ssDNA and the flexible C-terminal subdomain (CTD) of DRT4’s α-Rep domain. We further found that gp44 binding interferes with DRT4-mediated ssDNA synthesis. Importantly, both ssDNA binding and specific interactions with the CTD were required to activate DRT4’s RNase activity and confer antiphage defense. These findings establish the CTD as a critical determinant of DRT4 activation and reveal a dual requirement for nucleic acid binding and protein-protein recognition in licensing the immune response.

## Results

### DRT4 protects against diverse phages

DRT4 from *Escherichia coli* 21-C8-A (*Ec*DRT4), hereafter referred to as DRT4, has previously been shown to protect against phages T3, T5, T7, and φV-1 ^11,12,15^. Consistent with its reported activity against T5, DRT4-expressing cultures continued to grow following T5 infection at estimated multiplicities of infection (MOIs) of 0.1 and 1, but ceased growth at an MOI of 10 (**Supplementary Fig. 1A**). These population-level growth dynamics are consistent with abortive infection, in which infected cells arrest growth or die, limiting further phage propagation. In the absence of infection, DRT4-expressing cultures grew similarly to the empty-vector control, showing that DRT4 expression is not toxic under the conditions tested (**Supplementary Fig. 1B**). To systematically assess DRT4-mediated antiphage activity, we challenged *E. coli* MG1655ΔRM expressing DRT4 with a diverse panel of phages from the BASEL collection ^16^. DRT4 conferred broad protection against phages from multiple families (**Fig. 1B, Supplementary Fig. 2**).

To identify phage factors that trigger DRT4 activation, we isolated escape mutants of phage Bas17 (**Supplementary Fig. 3A**). Sequencing revealed mutations in the SSB-encoding gene *gp44* and an upstream DUF2815-associated non-coding RNA (ncRNA) (**Supplementary Fig. 3B**). DUF2815-associated ncRNAs have previously been proposed to act as feedback-regulatory elements, potentially mimicking ssDNA substrates to regulate translation of the associated SSB proteins ^17^. An AlphaFold 3 model predicted that gp44 comprises an N-terminal DUF2815 domain, which has been proposed to bind ssDNA ^18^, and an intrinsically disordered C-terminal tail (CT-Tail) containing four negatively and positively charged tandem repeats (**Supplementary Fig. 3C, D**). Escaper mutations included two individual substitutions in the DUF2815 domain, K72N and D152G; a 9-basepair (bp) duplication-insertion in the region encoding a negatively charged segment of the CT-tail; and a 16-bp deletion in the intergenic region upstream of *gp44*. One escape mutant carried both D152G and an ncRNA deletion (**Supplementary Fig. 3B**). The substituted gp44 residues, K72 and D152, were highly conserved among gp44 homologs (**Supplementary Fig. 3E**), suggesting a functional role, possibly for DNA binding.

The genomic context of *gp44* previously suggested a role in DNA metabolism ^17^. FoldSeek analysis revealed significant structural similarities between gp44’s DUF2815 and the replication protein ORF6 from *Enterobacter cancerogenus* phage ENC34 (**Supplementary Fig. 3F**). ORF6 adopts an oligosaccharide/oligonucleotide-binding (OB)-fold, a versatile structural motif implicated in interactions with nucleic acids, but also other proteins ^19,20^. This OB-fold similarity supports a putative nucleic acid-binding function for gp44’s DUF2815 domain and raises the possibility that gp44 also engages in protein interactions.

We identified homologs of gp44 only in *Dhillonviruse*s and *Guernseyvirinae*, but not in other phages susceptible to the DRT4-mediated immune response (**Fig. 1B, Supplementary Fig. 2**). DRT4 may therefore respond to other factors encoded by these phages. Previous reports that DRT4 is activated by T5 gp58 (ORF55) and T7 SSB support the possibility that other phage SSB proteins also trigger DRT4-mediated defense ^11,12^.

Together, these results demonstrate that DTR4 protects against diverse bacteriophages and further implicate phage SSB as triggers of DRT4.

### DRT4 assembles into a homohexameric complex

To investigate the molecular architecture of DRT4, we heterologously expressed, purified and determined its cryo-EM structure at an overall resolution of ∼2.2 Å (**Fig. 2A, Supplementary Figs. 4** and **5, and Supplementary Tab. 4**). The structure revealed that DRT4 comprises an N-terminal RT-like domain fused to a C-terminal α-Rep domain and assembles into a homohexamer with C3 symmetry (**Fig. 2A)**. In this complex, trimers stack back-to-back, with a relative rotation of approximately 10°, enclosing a central cavity. Residues 508-540 were not resolved in the cryo-EM map, indicating structural flexibility or proteolytic degradation of this C-terminal domain (CTD). Recovery of the protein through a C-terminal affinity tag during cryo-EM sample preparation favors flexibility as the explanation.

**Fig. 2.**
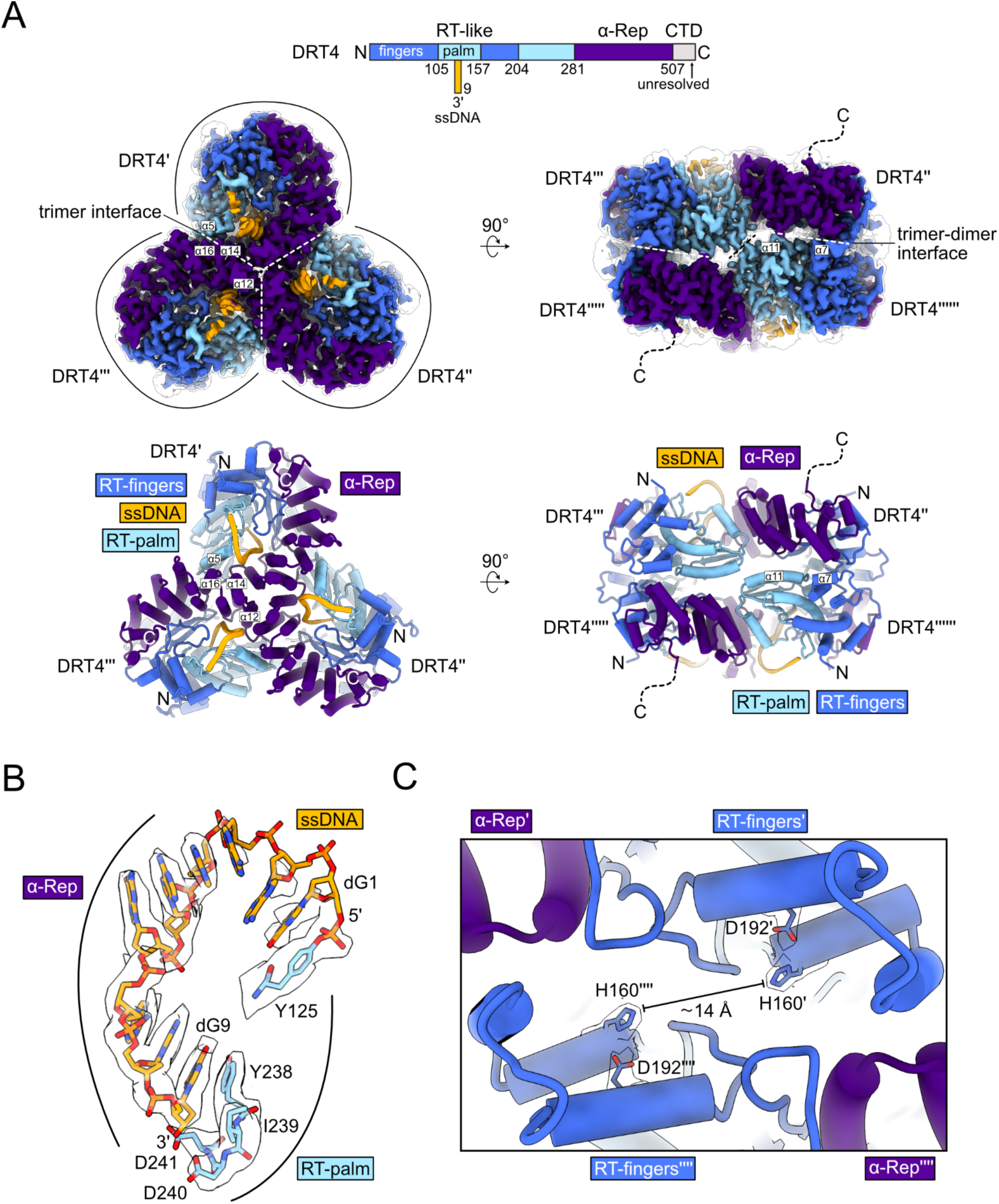
DRT4 assembles into a homohexameric complex. A, Cryo-EM maps (top) and corresponding model (bottom) of DRT4, shown in 90° rotations. Coloring according to the schematic above. C-shaped outlines highlight individual protomers. Dashed lines indicate interfaces within the complex. The unfiltered map is shown as a translucent surface. **B,** Close-up of the covalently attached ssDNA (orange) and interacting residues of the RT-palm subdomain (light blue) within the RT active site. The sharpened cryo-EM map is shown as a translucent surface. **C,** Close-up of the RNase active site, with active-site residues shown as sticks and the sharpened cryo-EM map displayed as a translucent surface.

Analysis of the subunit contacts identified three interfaces contributing to hexamer assembly (**Fig. 2A**). At the center of each trimer, α-helices 12 and 14 of the α-Rep domains form a trimerization interface. Additional contacts involve α-helix 5 of the RT palm subdomain and α-helix 16 of the α-Rep domain. At the interface between the two trimers, α-helices 7 and 11 of the RT fingers and palm subdomains pack against the α-Rep domains and RT palm subdomains of the opposing trimer.

Each subunit contained an ∼9 nucleotide ssDNA product covalently linked at its 5’-end to the surface-exposed tyrosine 125 (**Fig. 2B**). The ssDNA extended through a channel between the RT-palm and αRep domains towards the RT active site and the central cavity. Its 3’-terminal base was stabilized by stacking against tyrosine 238 of the catalytic YxDD RT-motif (**Fig. 2B**).

Overall, these structural features are consistent with previously reported structures of DRT4 and the related DRT6 and AbiK complexes ^8,10–12^.

### DRT4 adopts a polymerase-active, RNase-inactive conformation

We next examined the conformations of the RT and effector RNase active sites. Within the RT palm subdomain, the conserved YxDD catalytic motif and surrounding residues adopted a configuration consistent with polymerase activity (**Fig. 2B**). The ssDNA primer’s 3’-terminal base stacked against Y238, orienting the deoxyribose 3’-OH group toward the catalytic aspartates 240 and 241 of the YxDD motif (**Fig. 2B**).

The three surface-exposed RNase active sites lie at the interface between the two trimers, where the RT fingers and α-Rep domains are juxtaposed. Each site contained the catalytic HD motif formed by histidine 160 and aspartate 192 (**Fig. 2C**). The H160 residues contributed by opposing subunits were placed apart by ∼14 Å, closely matching the ∼15 Å distance reported for the RNase-inactive state ^12^. By contrast, the proposed RNase-active state shortens the distance of the histidines to ∼7 Å (**Fig. 2C and Supplementary Fig. 6**) ^12^.

These observations indicate that the structure captures DRT4 in a conformation consistent with an active polymerase and an inactive RNase, in agreement with previous reports.

### DRT4 synthesizes purine-rich ssDNA

Our cryo-EM structure revealed short oligonucleotides covalently linked to the hydroxyl group of Y125, consistent with protein-primed DNA synthesis. To determine their nucleotide composition, we purified these products and analyzed them by mass spectrometry. The products contained dAMP and dGMP at an approximate ratio of 2:1 (**Fig. 3A**). Mass spectrometry further identified deoxyguanosine as the nucleotide linked to Y125 (**Supplementary Fig. 7**), suggesting that dGTP serves as the initiating substrate for ssDNA synthesis.

**Fig. 3.**
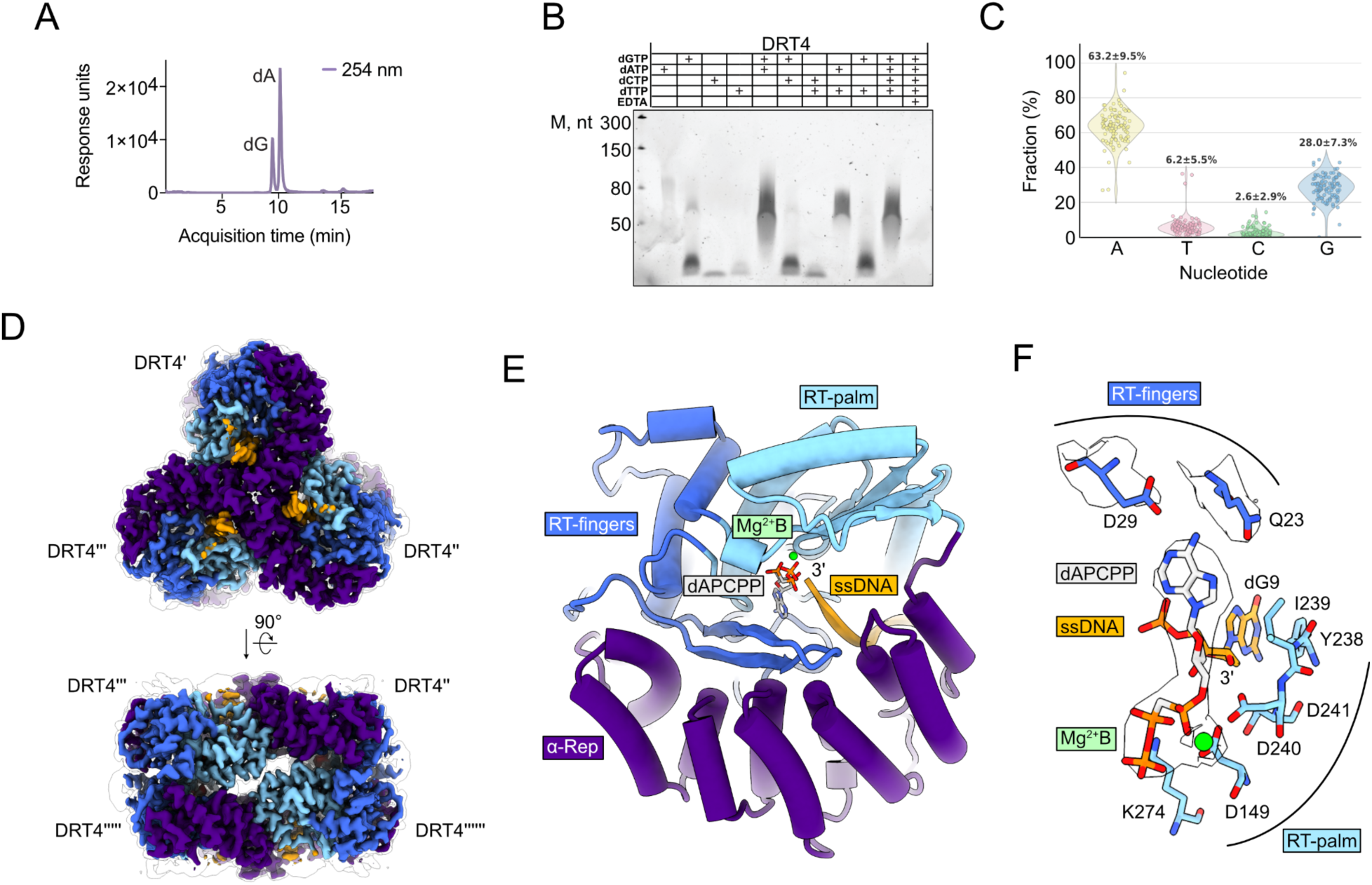
DRT4 synthesizes purine-rich ssDNA. A, HPLC-MS analysis of ssDNA synthesized *in vitro*. *n* = 1. **B,** Urea-PAGE analysis of ssDNA products synthesized with the indicated nucleotide substrates. Representative data; *n* ≥ 2 technical replicates. **C.** Nucleotide composition of DRT4-produced ssDNA determined by long-read sequencing. *n* = 1. **D,** Cryo-EM maps of hexameric DRT4 bound to the non-hydrolyzable dATP analogue dAPCPP. The unfiltered map is shown as a translucent surface, and the sharpened map is colored according to the schematic in Fig. 2A. **E,** Model of an individual DRT4 subunit viewed from the central cavity, colored as in panel D. Bound dAPCPP is shown as light-gray sticks. **F.** Close-up of the nucleotide-binding pocket showing dAPCPP (light-gray sticks), the magnesium cofactor (green sphere), and residues D29 and Q23 (dark-blue sticks). The surrounding sharpened cryo-EM map is shown as a translucent surface.

We next examined DRT4’s nucleotide preference during ssDNA synthesis *in vitro*. We incubated DRT4 with all four dNTPs in the presence of Mg^2+^, a physiologically relevant divalent cation commonly used as a cofactor by enzymes involved in nucleotide and nucleic acid metabolism ^21–23^. DRT4 synthesized ∼50-100 nucleotide long ssDNA products and preferentially incorporated dATP and dGTP (**Fig. 3B**). Nanopore sequencing confirmed that the products predominantly contained deoxyadenosine and deoxyguanosine at an approximate ratio of 2:1 (**Fig. 3C**), consistent with the mass spectrometry results (**Fig. 3A**). Multiple sequence alignment revealed no apparent shared sequence pattern, suggesting that DRT4 synthesizes purine-rich ssDNA without a defined sequence (**Supplementary Fig. 8**).

Together, these results demonstrate that DRT4 preferentially incorporates purine deoxynucleotides during Mg^2+^-dependent ssDNA synthesis, consistent with a recent report ^11^.

### Q23 and D29 contribute to deoxynucleotide substrates positioning

To investigate the structural basis of DRT4’s preference for dATP, we incubated DRT4 with the non-hydrolysable dATP analogue dAPCPP and determined the cryo-EM structure at an overall resolution of ∼2.4 Å (**Fig. 3D and E, Supplementary Fig. 9, Supplementary Tab. 4**). The resulting map revealed segments for dAPCPP and a Mg^2+^ ion near the YxDD motif in the RT active sites (**Fig. 3F**).

The Mg^2+^ ion occupied metal-binding site B, where it was coordinated by D149 and positioned to interact with the triphosphate moiety of the incoming nucleotide (**Fig. 3F**). No density was observed for a putative second Mg^2+^ ion at site A, where coordination by D240 and D241 could facilitate activation of the ssDNA primer’s 3’-OH group for nucleophilic attack on the nucleotide’s α-phosphate. These observations are consistent with the structure capturing a pre-catalytic state of DRT4.

Within the RT fingers subdomain, the side chains of glutamine 23 and aspartate 29 projected into the nucleotide-binding pocket and were positioned to contact the adenine base of dAPCPP (**Fig. 3F**). These contacts may contribute to DRT4’s preference for purine nucleotides.

For dGTP, an alternative rotamer of the Q23 side chain could potentially allow its amide nitrogen (Nε2) to hydrogen-bond with the O6 carbonyl group of guanine. By contrast, the smaller pyrimidine bases of dCTP and dTTP may be unable to form equivalent interactions with Q23 or D29.

These observations suggest that Q23 and D29 contribute to nucleotide selection by DRT4 and provide a possible explanation for its preferential incorporation of dATP and dGTP.

### gp44 is predicted to interact with DRT4’s CTD and ssDNA

To investigate the interaction between DRT4 and gp44, we cloned the *gp44* gene and purified the protein (**Supplementary Fig. 10A**). Initial attempts to clone wild-type *gp44* were unsuccessful, potentially owing to toxicity associated with its expression. Cloning was successful when the predicted upstream ncRNA sequence was included in the construct. Purified gp44 had an A_260_/A_280_ ratio of ∼0.5, consistent with no or minimal nucleic acid contamination.

*In vitro* reconstitution of DRT4-gp44 complexes revealed that gp44 bound to DRT4, irrespective of the extended ssDNA being produced by DRT4 (**Fig 4A**). We next analyzed the reconstituted DRT4-gp44 complexes by cryo-EM (**Supplementary Fig. 11**). However, reconstruction yielded only a low-resolution map, precluding detailed structural analysis (**Supplementary Fig. 11B**). This result was consistent with gp44-induced destabilization or increased structural heterogeneity of DRT4 under the tested conditions.

**Fig. 4.**
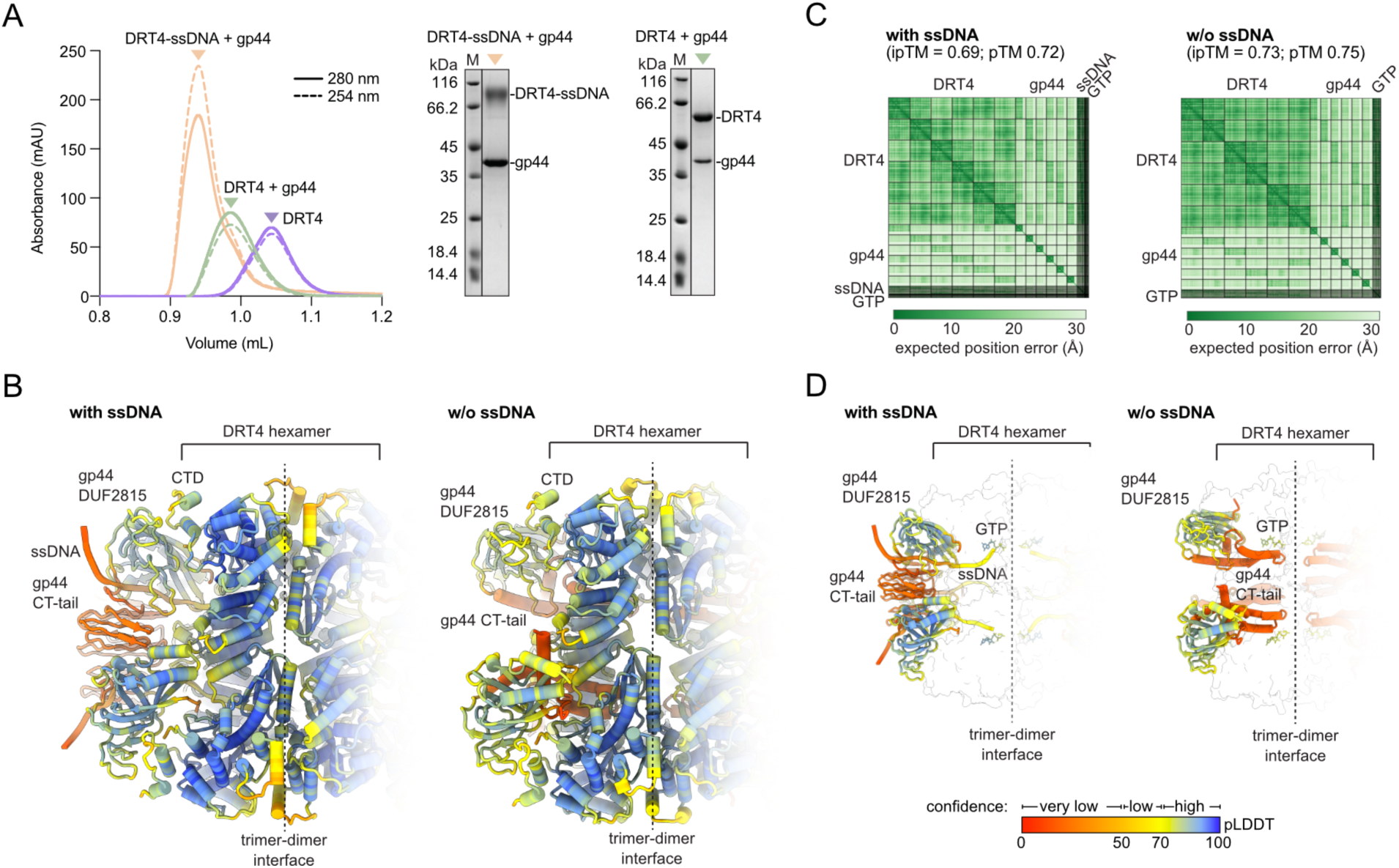
Reconstitution and structure prediction of the DRT4-gp44 complex. **A**, *In vitro* reconstitution of DRT4 and gp44. Left, size-exclusion chromatography traces; colored triangles indicate peak fractions analyzed by SDS-PAGE (right). Representative data; *n* ≥ 2 technical replicates. **B,** AlphaFold 3 models of DRT4 bound to nucleotides and gp44 in the presence (left) or absence (right) of ssDNA. **C,** Predicted aligned error (PAE) matrices for the models shown in B. **D,** Predicted conformations of gp44 in the presence (left) or absence (right) of ssDNA, with the DRT4 hexamer shown as a translucent surface. Models in panels B and D are colored by predicted local distance difference test (pLDDT) scores according to the rainbow scale in D.

**Fig. 5.**
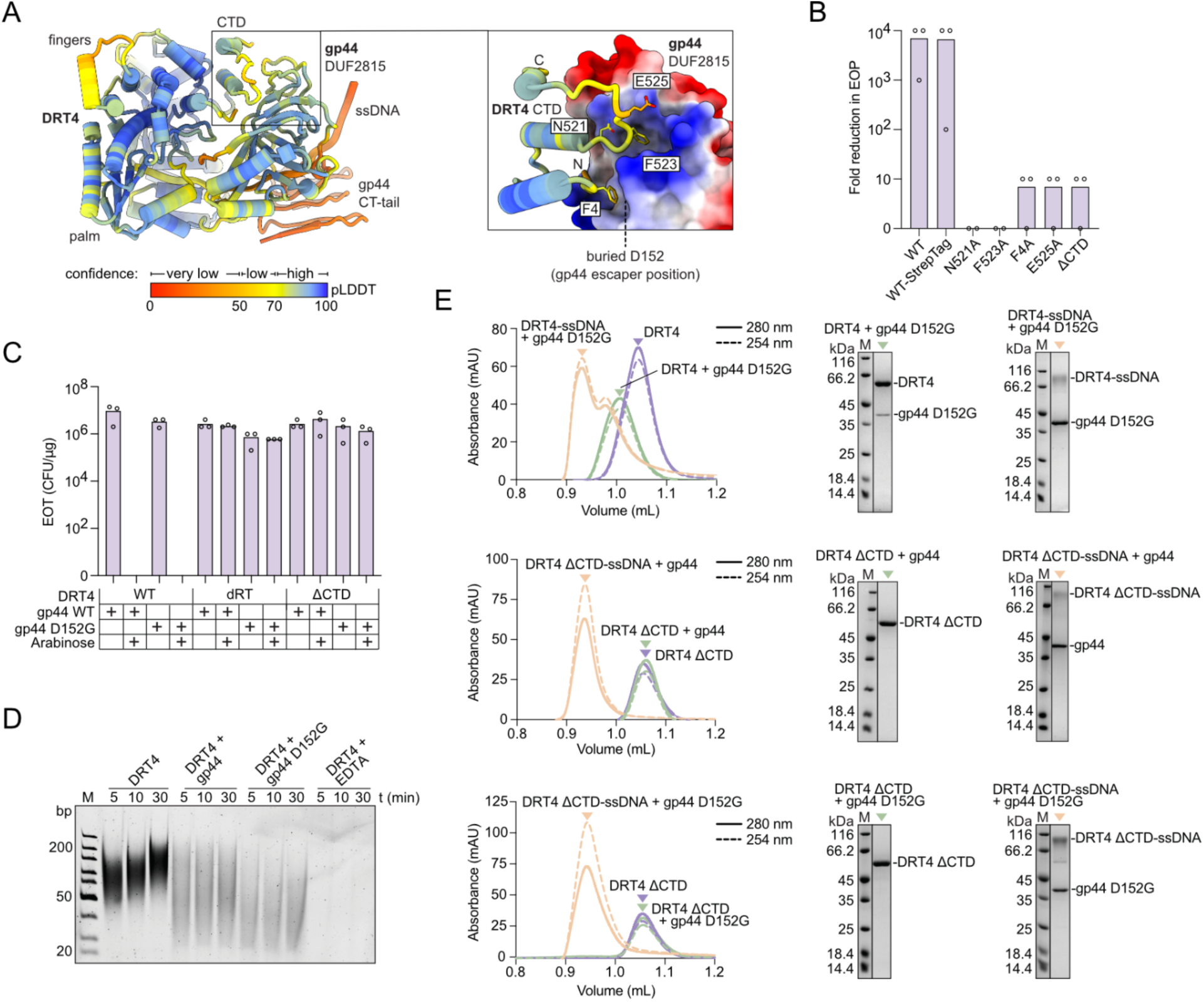
The DRT4 CTD-gp44 interface is essential for antiphage defense. A, Left, predicted gp44 and DRT4-ssDNA heterodimer (from dodecamer model shown in Fig. 4B), colored by pLDDT score. Right, close-up of the predicted interface between the DRT4 CTD and the gp44 DUF2815 domain. DRT4 is colored by pLDDT, and gp44 is shown as a surface colored by electrostatic potential. **B,** Antiphage activity of DRT4 variants against Bas17, measured by efficiency of plating (EOP). *n* = 3 technical replicates; mean. **C,** Efficiency of transformation (EOT) of plasmids encoding gp44 or its variants into cells expressing DRT4 or its variants. *n* = 3 technical replicates; mean. **D,** PAGE analysis of ssDNA products synthesized by DRT4 in the presence of wild-type gp44 or the D152G escape variant. Representative data; *n* ≥ 2 technical replicates. **E,** *In vitro* complex reconstitution of DRT4 and gp44 variants. Left, size-exclusion chromatography traces; colored triangles indicate peak fractions analyzed by SDS-PAGE (right). Representative data; *n* ≥ 2 technical replicates.

To nonetheless explore the molecular basis of gp44-dependent activation, we predicted the DRT4-gp44 complex using AlphaFold 3 (**Fig. 4B**). The predictions included six molecules each of DRT4 and gp44, Mg^2+^ ions, and GTP as a proxy for dGTP. To assess the potential contribution of ssDNA, we also generated a model containing six ssDNA molecules composed of dA and dG. The predicted model with ssDNA yielded ipTM and pTM scores of 0.69 and 0.72, respectively, whereas the prediction without ssDNA yielded corresponding scores of 0.73 and 0.75 (**Fig. 4C**). These scores supported a reasonable overall confidence of the predicted complex models but did not establish the accuracy of individual contacts.

Both predicted models retained the overall hexameric architecture of our experimentally determined DRT4 structures, while additionally suggesting gp44 interactions. In both models, the gp44 DUF2815 domains were positioned near the DRT4 CTDs; in the ssDNA-containing model, the ssDNAs were placed within DRT4’s DNA binding channel and associated with gp44 DUF2815 domains (**Fig. 4B**). The gp44 CT-tails were modeled with low confidence in both predictions (**Fig. 4D**). In the presence of ssDNA, they bundled-up over the center of the trimer surface between the three gp44 DUF2815 domains. In the absence of ssDNA, they were predicted to occupy the DNA-binding channels of DRT4.

Together, these predictions suggest that the gp44 DUF2815 domain may directly contact both the DRT4 CTD and ssDNA. The predicted placement of the gp44 CT-tail within the DNA-binding channel further raises the possibility that it competes with ssDNA for channel occupancy, potentially promoting DNA displacement. Importantly, this proposed role remains tentative given the low confidence of the CT-tail models and the lack of experimental data.

### The DRT4 CTD and RT catalytic activity are required for gp44-dependent activation

To investigate the predicted DRT4-gp44 interface, we identified candidate DRT4 residues that could contribute to gp44 binding (**Fig. 5A**). The N-terminal residue phenylalanine 4 and CTD residues asparagine 521, phenylalanine 523, and glutamate 525 were positioned at the predicted interface near the central β-sheet of gp44’s DUF2815 domain. We generated individual alanine substitutions at these positions (F4A, N521A, F523A, and E525A) and a CTD deletion variant (ΔCTD; residues 508-540). All variants lost the ability to protect against phage Bas17 *in vivo* (**Fig. 5B**).

To assess whether this loss of protection reflected defects in protein production, solubility, or ssDNA synthesis, we purified each variant and measured its polymerase activity *in vitro*. All variants were recovered as soluble proteins and retained ssDNA synthesis activity comparable to that of wild-type DRT4 (**Supplementary Figs. 12 and 13**). Thus, disruption of the predicted interface abolished antiphage protection without impairing ssDNA synthesis. Expression of DRT4 ΔCTD also did not measurably impair bacterial growth, suggesting that the deletion is not toxic under the tested conditions (**Supplementary Fig. 14A**).

We next examined whether the CTD contributes to gp44-dependent DRT4 activation. In efficiency-of-transformation (EOT) assays, introduction of a construct encoding wild-type ncRNA-gp44 reduced transformation efficiency in cells expressing wild-type DRT4, but not in cells expressing the catalytically inactive RT mutant (dRT) or DRT4 ΔCTD (**Fig. 5C**). Expression of ncRNA-gp44 in the absence of DRT4 did not impair bacterial growth, supporting a DRT4-dependent phenotype (**Supplementary Fig. 14B,C**).

Interestingly, the gp44 D152G escaper variant also reduced EOT in cells expressing wild-type DRT4 (**Fig. 5C**), despite carrying a substitution near the predicted CTD-binding interface (**Fig. 5A**). One possible explanation is that elevated expression in the transformation assay compensates for an impaired interaction with DRT4 that limits activation at native expression levels during infection.

Together, these results identify the CTD and RT catalytic activity as requirements for gp44-dependent DRT4 activation. Combined with the structural predictions, they support a model in which activation depends on both a specific DRT4-gp44 interaction and DRT4-mediated ssDNA synthesis.

### gp44 associates with DRT4 through CTD- and ssDNA-dependent interactions

Incubation of DRT4 with gp44 in the presence of dNTPs yielded shorter ssDNA products than reactions lacking gp44, indicating that gp44 alters DRT4-mediated ssDNA synthesis (**Fig. 5D**). This observation is consistent with gp44 interacting with DRT4 or its ssDNA product.

To examine the basis of this interaction, we performed *in vitro* reconstitution experiments with DRT4 and either wild-type gp44 or the gp44 D152G escaper variant, with or without dNTPs (**Fig. 5E**). As shown above, in the presence of dNTPs, DRT4 synthesized ssDNA, whereas without added dNTPs, covalently attached short ssDNA products were retained (**Figs. 2 and 3**). Wild-type DRT4 co-eluted with both gp44 variants under all conditions tested (**Fig. 5E**), indicating that complex formation does not require ongoing ssDNA synthesis.

To assess the contribution of the CTD, we repeated these experiments using DRT4 ΔCTD. This variant co-eluted with gp44 only in the presence of dNTPs, when ssDNA synthesis occurred (**Fig. 5E**). Thus, the CTD was required for detectable association in the absence of ongoing synthesis, whereas newly synthesized ssDNA supported complex formation without the CTD.

These findings support a model in which gp44 associates with DRT4 through contacts with both its CTD and its ssDNA product.

### The DRT4 CTD is required for gp44-triggered RNase activation

We next tested whether gp44 and deoxynucleotides stimulate DRT4’s RNase activity using a fluorophore-quencher RNA reporter assay. Reporter cleavage was significantly accelerated only when gp44 was combined with either dGTP alone or a mixture of all four dNTPs, while other individual dNTPs did not produce comparable stimulation (**Fig. 6A, Supplementary Fig. 15A**).

**Fig. 6.**
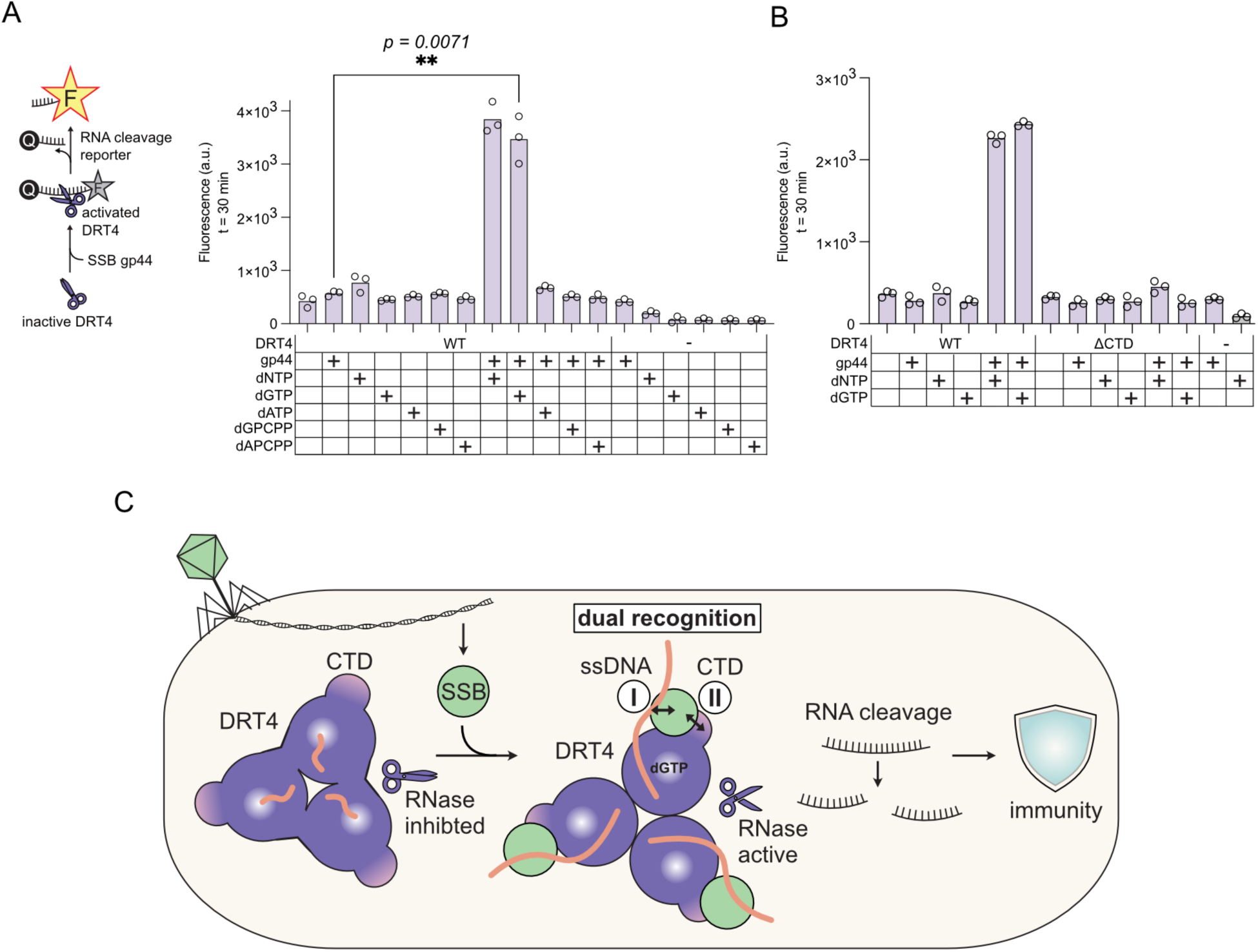
The DRT4 CTD is required for full gp44-dependent RNase activation. A, Left, schematic of the fluorophore-quencher (FQ) RNA reporter assay. Right, DRT4 RNase activity measured using the FQ-reporter in presence of gp44 and indicated nucleotides. *n* = 3 technical replicates; mean, student’s t-test. **B,** FQ-reporter assay measuring RNase activity of DRT4 ΔCTD relative to wild-type DRT4. *n* = 3 technical replicates; mean. **C,** Proposed model of DRT4 activation via a dual SSB recognition mechanism. White circles indicate the two requirements for SSB gp44 recognition: (I) ssDNA binding, and (II) specific protein-protein interactions involving the DRT4 CTD to license RNase activation for antiphage immunity.

This shows that gp44-dependent RNase activation requires dGTP under the conditions tested, consistent with previous observations for T7 SSB ^12^. These results suggest that distinct phage SSB proteins activate DRT4 through a shared mechanism.

To determine whether this activation requires the CTD, we repeated the assay with DRT4 ΔCTD. Unlike wild-type DRT4, the ΔCTD variant failed to support rapid reporter cleavage in the presence of gp44 and either dGTP alone or all four dNTPs (**Fig. 6B, Supplementary Fig. 15B**). This shows that the CTD is required for full gp44-dependent stimulation of DRT4’s RNase activity.

Together with the complex reconstitution and ssDNA synthesis assays, these results support a model in which full RNase activation requires gp44 engagement with both DRT4-produced ssDNA and the CTD.

## Discussion

Our study defines the structural and functional requirements for activation of DRT4 by the phage SSB protein gp44. DRT4 protects against diverse bacteriophages, preferentially synthesizes purine-rich ssDNA, and requires its sensory CTD domain for gp44-dependent RNase activation. These findings support a “two-factor authentication” model in which DRT4-produced ssDNA recruits phage SSB, while CTD-mediated recognition licenses RNase activation (**Fig. 6C**).

Protection against multiple *E. coli* phage families suggests that DRT4 recognizes features shared by diverse phages, although the activating factors in several susceptible phages remain unidentified. Recognition of replication-associated proteins such as SSBs could extend protection beyond closely related viruses ^24^. DRT4-expressing cultures continued to grow following T5 infection at lower multiplicities of infection but underwent growth arrest at a higher multiplicity. Together with previous reports of DRT4-dependent cell death during T5 and T7 infection, these observations support an abortive infection mechanism that limits phage propagation through growth arrest or death of infected cells under the tested conditions ^11,12,25^ .

Our structures revealed the conserved homohexameric architecture of DRT4 and covalently attached ssDNA products consistent with protein-primed synthesis ^11,12^. The dAPCPP-bound structure places Q23 and D29 near the nucleotide base, suggesting a molecular basis for purine selection. Consistently, DRT4 predominantly synthesized dA/dG-containing ssDNA under Mg^2+^-dependent conditions. Previous observations of less specific deoxynucleotide incorporation in the presence of Mn^2+^ suggest that metal identity influences substrate discrimination ^11^, in agreement with the generally greater selectivity supported by Mg^2+^ ^23,26,27^. Purine-rich ssDNA is therefore a plausible physiological product, given the high concentration of Mg^2+^ in cells ^22^. However, the ssDNA-composition during infection remains to be determined.

The CTD of DRT4 provides a previously unrecognized direct link between gp44 association and RNase activation. Its deletion abolished protection and prevented full RNase stimulation. The CTD also supported gp44 association without ongoing DNA synthesis, whereas newly synthesized ssDNA supported gp44 association in the ΔCTD variant. These results extend the proposed requirement for ssDNA, phage SSB, and dGTP in the activation of DRT4’s RNase, by identifying an additional CTD-dependent checkpoint. However, the precise molecular interactions and conformational dynamics coupling gp44 binding to RNase activation remain unresolved, and the predicted displacement of ssDNA by the gp44 CT-tail will require future experimental validation.

Increased dNTP availability during infection has been proposed to promote DRT4 activation in concert with the triggering by phage SSB ^12^. Expression of gp44 resulted in a DRT4-dependent plasmid transformation defect without infection, indicating that other infection-associated changes were not required for DRT4 activation under the tested conditions. Elevated dNTP levels during phage infection may therefore not necessarily be a critical determinant of DRT4 activation.

Phage SSBs activate diverse bacterial defense systems, emphasizing these proteins involved in replication, recombination, and repair as recurring signals of infection ^8,14,24,28–35^. Although these systems share SSBs as common activation signals, their activation mechanisms differ. While some are activated indirectly ^28–30,33,35^, others may sense SSBs by direct protein-protein interactions ^31,34^. DRT4 appears to combine recognition of SSB’s ssDNA-binding activity with specific CTD-mediated protein contacts. This dual requirement could confer selectivity and potentially allow DRT4 to distinguish phage SSBs from their host counterparts. The requirement for C-terminal residues in AbiK-mediated defense further suggests a broader functional role for these regions in related defense-associated RTs ^36^.

In summary, our findings reveal how DNA synthesis and direct protein recognition can cooperate to control an abortive infection phenotype. Our model places ssDNA-mediated SSB recruitment and CTD-dependent recognition as complementary requirements for RNase activation. Such a two-factor mechanism could coordinate broad detection of viral proteins with a selective immune response, illustrating how bacterial defenses integrate molecular cues before committing to growth arrest or cell death.

## Materials and Methods

### BASEL phage plaque assay screen

*E. coli* MG1655ΔRM ^37^ were transformed either with the empty control vector pACYC184 (NEB) or the DRT4-carrying plasmid pLG012 ^15,37^ (Addgene ID:157890) and plated on LB-agar-chloramphenicol (10 g/L NaCl, 10 g/L tryptone, 5 g/L yeast extract, agar 15 g/L, Cm 25 µg/mL), prior to incubation overnight at 37 °C. 4 mL LB-Cm (10 g/L NaCl, 10 g/L tryptone, 5 g/L yeast extract, Cm 25 µg/mL) cultures were subsequently inoculated and incubated for 16 h at 37 °C to prepare overnight cultures for double-layer agar plaque assays. Double-layer agar plates were prepared using 1.5% agar LB-Cm (Cm, 25 µg/mL) for the bottom layer and LB-Cm (Cm, 25 µg/mL) 0.75% agar mixed with 200 µL of overnight *E. coli* culture for the top layer. For effective infection, 20 mM MgCl_2_ and 5 mM CaCl_2_ were supplemented into the LB-agar. 10-fold serial phage dilutions were spot-plated in 3.5 µL volumes onto the double-layer plates, prior to incubation at 37 °C for 16 h.

### Isolation of Bas17 phage mutants

Escaper phages were isolated as previously described ^38^. In brief, to generate DRT4-escaper phages, *E. coli* MG1655ΔRM were transformed with plasmid pLG012 ^15^ (Addgene ID:157890), plated on LB-Agar plates containing chloramphenicol (Cm, 25 µg/mL) and incubated for 16 h at 37 °C. 4 mL LB-Cm cultures were subsequently inoculated and incubated for 16 h at 37 °C to prepare overnight cultures for double-layer agar plaque assays. Double-layer agar plates were prepared by layering 30 mL 1.5% LB-agar-Cm for the bottom layer and 6 mL 0.75% LB-agar-Cm mixed with 500 µL overnight culture for the top layer. Subsequently, 10-fold dilutions series of Bas17 were spot plated in 3.5 µL volumes onto the top layer, prior to incubation for 16 h at 37 °C. Emerging escaper phage candidates, appearing to propagate well in presence of DRT4 as indicated by large plaques at high dilution, were subsequently isolated by scraping spots for transfer into 1 mL phage buffer (50 mM Tris pH 7.4, 10 mM NaCl, 100 mM MgCl_2_). Isolated phages were incubated for 1 h at room temperature prior to repeated vortexing for phage release. The agar was then removed by centrifugation at 3,200 x g for 10 min and the phage supernatant was transferred to a new tube. Isolated phages were then replated as serial dilutions on fresh DRT4-expressing *E. coli* MG1655ΔRM double layer plates (LB-agar-Cm) and incubated for 16 h at 37 °C. Enriched escaper phage spots were subsequently used for the final propagation and purification step, where individual plaques were transferred to 4 mL LB-Cm containing *E. coli* MG1655ΔRM at an OD_600_ ≈ 0.5. Phages were subsequently incubated at 37 °C, shaking at 250 r.p.m. for ∼20 h. To separate phage from bacteria, cells were sedimented by centrifugation at 3,200 x g for 5 min. The phage supernatant was then filtered using a 0.22 μm filter and stored at 4 °C.

### Phage DNA extraction

Prepared phage escaper solutions were propagated on a 10 mL *E. coli* MG1655ΔRM culture in LB-Cm (Cm, 30 µg/mL) at 37 °C overnight. The next day, cultures were centrifuged for 5 min at 3,200 x g, and the supernatant was filtered through a 0.22 μm pore filter to remove cell debris and intact cells. Filtered supernatants were centrifuged at 12,000 g for 1 h, the supernatant was discarded and phages were resuspended in 400 µL of 30 mM Tris-HCl pH 7.5, 50 mM MgCl_2_, 0.5 mM CaCl_2_, prior to incubation overnight at 4 °C. 1 µL of RNase A (10 mg/mL) and 5 µL of DNase I (1 U/µL) were added the next day and samples were incubated at 37 °C for 2 h. Subsequently, EDTA (final concentration of 62.5 mM) and 15 µL of Proteinase K (0.65 mg/mL) were added to the mix and incubated at 56 °C for 2 h. Phage genomic DNA was purified using the phenol/chloroform extraction, adding 400 μL of phenol/chloroform/isoamyl alcohol (ratio 25:24:1; Roth #A156.2) to aqueous fraction, with 3 phenol/chloroform extractions and 2 chloroform wash steps. The nucleic acids phase was supplemented with 40 μl volume of 3 M sodium acetate pH 5.2 and 1 mL volume of −20 °C ethanol. Samples were mixed and precipitated overnight at −20 °C. DNA was collected by centrifugation for 60 min, 12,000 g, at 4 °C and the supernatant was removed. Pellets were washed 3 times with cold −20 °C 70% ethanol and centrifuged for 5 min, 12,000 g, at 4 °C after each wash step. Subsequently DNA pellets were air dried. DNA pellets were dissolved in water and the concentration of nucleic acids was estimated based on the absorbance at 260 nm using a NanoDrop Eight spectrophotometer (Thermo Scientific).

### Phage genome sequencing and escaper analysis

Phage genomic DNA sequencing libraries were prepared using the Colibri ES DNA Library Prep Kit (Invitrogen) following the manufacturer’s instructions and sequenced on an Illumina MiniSeq in 150 PE mode (MiniSeq Control Software v2.2.1.9; LRM Version v2.4.1.2636; RTA Version v2.11.4.0; Module Version Title 2.0.1.17).

Sequencing data quality was evaluated using FastQC (v0.12.1; https://www.bioinformatics.babraham.ac.uk/projects/fastqc/) and MultiQC^39^ (software version 1.30). Adapters were trimmed with BBDuk from the BBMap package (software version 38.18; https://sourceforge.net/projects/bbmap; ktrim = r, *k* = 15, mink = 11, hdist = 1, tpe, tbo). Further analysis was performed using breseq^40^ (software version 0.37.1). As breseq requires coverage of ∼100, the reads were subsampled to 40,000 per dataset using seqtk –s 123 (v1.5; https://github.com/lh3/seqtk). The wild-type Bas17 phage genome (GenBank accession: <u>NC_073057.1)</u> was used as a reference, and analysis was performed using breseq with default parameters. The output was evaluated and summarised manually.

To identify gp44 homologs across BASEL phage collection, 3 iterations of PSI-BLAST against the NCBI non-redundant protein sequence (nr) database were performed (query GenBank accession: <u>QXV81960.1</u>). Sequences exhibiting 30–90% sequence homology and >70% query coverage (expected threshold: 0.05, BLOSSUM62 substitution matrix) were selected. The resulting sequences were clustered to 80% sequence identity and 80% alignment coverage using the MMseqs2 algorithm. Sequences were aligned using European Bioinformatics Institute (EMBL-EBI) multiple sequence alignment (MSA) tool with the default MAFFT algorithm (gap opening penalty: 1.53). The Hidden Markov model (HMM) profiles were built using the hmmbuild command from HMMER software version 3.4 (http://hmmer.org/). The resulting HMM profiles were used to search the proteomes with the hmmsearch command E-value:0.001; http://hmmer.org/.

### Plasmid construction

Constructs for expression and affinity purification for DRT4 were cloned by fusing a Strep-tag II encoding sequence downstream of the *drt4* gene encoded on pLG012 ^15^ (Addgene plasmid number: #157890). Mutations encoding amino-acid substitutions of *drt4* variants were introduced by Golden Gate(GG)-mutagenesis.

Constructs for expression and affinity purification for gp44 and its escaper variant were fused to a Strep-tag II encoding sequence downstream the *ncRNA-gp44* genes encoded on Bas17 genomic DNA and cloned into pRSFDuet-1 using GG-assembly.

For EOT assays and toxicity evaluation, constructs encoding *ncRNA*-*gp44* variants were produced by fusing a His-tag-encoding sequence downstream of the respective *gp44* genes and cloned into pBAD using GG-assembly.

### *E. coli* growth assay

For DRT4 toxicity evaluation, *E. coli* MG1655 were transformed either with the empty control vector pACYC184 (NEB), or the plasmid pMJ016, containing a Strep-tag II encoding sequence fused downstream to the *drt4* gene, and plated on LB-agar-Cm (Cm, 25 µg/mL), prior to incubation overnight at 37 °C.

For gp44 toxicity evaluation, *E. coli* MG1655 were transformed either with the empty control vector pBAD, or the *ncRNA-gp44* plasmids pMJ024 and pMJ025, containing a C-terminal His tag II, and plated on LB-agar-Amp (Amp, 50 µg/mL), prior to incubation overnight at 37 °C. 4 mL LB-Cm or LB-Amp cultures were subsequently inoculated and incubated for 16 h at 37 °C to prepare overnight starter cultures. The next day, 4 mL LB-Cm (25 µg/mL Cm) or LB-Amp (50 µg/mL) media were inoculated with starter cultures and incubated at 37 °C, 200 r.p.m. until tOD_600_ ≈ 0.7. Subsequently, cultures were diluted to an OD_600_ ≈ 0.11 in 90 µL LB-Cm (25 µg/mL) or LB-Amp (50 µg/mL) and transferred into clear 96-well PS F-bottom plates (Greiner). Phages, or medium, were then added to a total volume of 100 µL and final OD_600_ of 0.1. Expression of gp44 was induced with 0.2 % arabinose. Phage titers were estimated by spot plating prior to the experiment.

Growth was monitored every 5 min for 16 h by OD-measurement in a CLARIOstar plus plate reader (BMG Labtech, software version 5.70 R2), at 37 °C, 200 r.p.m.. Data were analyzed and plotted in Prism (software version 10.6.1 (799), Graphpad).

### Protein production and purification from *E. coli* cultures

All proteins were purified at a constant temperature of 4 °C and kept on ice.

DRT4-encoding plasmids were transformed into *E. coli* DH5α (Thermo Scientific) plated on LB-agar plates containing chloramphenicol (Cm, 25 μg/mL) and incubated at 37 °C overnight. To express DRT4, 4 L TB-Cm (12 g/l tryptone, 24 g/l yeast extract, 4 mL/l glycerol, 17 mM KH_2_PO_4_, 72 mM K_2_HPO_4,_ Cm 25 μg/mL) were directly inoculated from transformation plates with multiple colonies and incubated at 37 °C shaking at 200 r.p.m. for 16 h. Cells were subsequently harvested and resuspended in a 40 mL lysis buffer #1 (100 mM Tris-HCl, pH 8.0, 150 mM NaCl, 1 mM EDTA) prior to cell disruption using an LM10 Microfluidizer (Microfluidics). Lysates were clarified by centrifugation at 20,000 x g for 40 min at 4 °C. The supernatants were subsequently loaded onto 1 mL StrepTrap XP columns (Cytiva) pre-equilibrated in lysis buffer #1 and washed with 20 column volumes (CV) of 100 mM Tris-HCl, pH 8.0, 150 mM NaCl, 1 mM EDTA buffer prior to elution with 6 CV elution buffer #1 (100 mM Tris-HCl, pH 8.0, 150 mM NaCl, 1 mM EDTA, 50 mM biotin). The elution was then applied to a 5 mL HiTrap Q HP column (Cytiva), pre-equilibrated in a buffer containing 20 mM NaCl and 10 mM Tris-HCl (pH 8.0). After a 4 CV column wash with 20 mM NaCl and 10 mM Tris-HCl (pH 8.0) buffer, a 15 CV gradient from 10 mM to 2 M NaCl in 10 mM Tris-HCl (pH 8.0) was applied for elution. The peak fraction was concentrated and further purified by SEC using a Superose 6 10/300 (Cytiva) column pre-equilibrated in SEC buffer #1 (100 mM Tris-HCl, pH 8.0, 150 mM NaCl). Peak fractions were concentrated to approximately 100 µL and concentrations were estimated based on the absorbance at 280 nm using a NanoDrop Eight spectrophotometer (Thermo Scientific). To express DRT4 ΔCTD, 0.6 L TB-Cm (Cm, 25 μg/mL) were directly inoculated from transformation plates with multiple colonies and incubated at 37 °C shaking at 200 r.p.m. for 16 h. Cells were subsequently harvested and resuspended in 75 mL lysis buffer #1 prior to cell disruption using an LM10 Microfluidizer (Microfluidics) equipped with an H10Z interaction chamber. Lysates were clarified by centrifugation at 20,000 x g for 20 min at 4 °C. Supernatants were loaded onto 600 µL Strep-Tactin XT resin (Cytiva) and incubated for 10 min on ice while shaking. Subsequently, the resin was centrifuged at 4,000 x g for 5 min at 4°C and the supernatant was discarded. The resin was then resuspended in 700 µL lysis buffer #1 and transferred to a Zymo-Spin P1 column (Zymo research). To remove non-specific contaminants, the resin was washed 3 times with 700 µL lysis buffer #1 prior to incubation with 600 µL elution buffer #1 for 5 min on ice. Proteins were eluted by centrifugation 4,000 x g for 1 min at 4 °C and concentrated to approximately 50 µL. The sample was further purified by SEC using Superose 6 3.2/300 (Cytiva) column, pre-equilibrated in SEC buffer #1. Peak fractions were concentrated to approximately 50 µL and concentration was estimated based on the absorbance at 280 nm using a NanoDrop Eight spectrophotometer (Thermo Scientific). The concentrated sample was split in aliquots, snap frozen in liquid nitrogen and stored at −70 °C prior to usage for assays.

For small-scale expression of His-tagged ncRNA-gp44 and its variants, pBAD plasmids encoding the *ncRNA*-*gp44* genes were transformed in in *E. coli* BL21 Star (DE3) (Invitrogen) and plated on LB-agar plates containing ampicillin (Amp, 50 µg/mL). To express the *ncRNA*-*gp44* genes, 4 mL LB-Amp overnight cultures, grown shaking at 37 °C, were used to inoculate 50 mL TB-Amp (Amp, 50 µg/mL) and incubated shaking at 200 r.p.m and 37 °C until OD_600_≈ 0.6. Gene expression was induced with 0.2 % arabinose (Carl Roth), prior to incubation for 16 h at 16 °C. Cells were subsequently collected by centrifugation and resuspended in a lysis buffer #2 (100 mM HEPES, pH 7.5, 1 M NaCl, 5 % glycerol, 40 mM imidazole) prior to cell disruption at 10,000 PSI using an LM10 Microfluidizer (Microfluidics). Lysates were clarified by centrifugation at 20,000 x g for 20 min at 4°C. Supernatants were loaded onto 100 µL HisPur Ni-NTA Resin (Thermo Fisher Scientific) and incubated for 10 min on ice while shaking. Subsequently, the resin was centrifuged at 4,000 x g for 5 min at 4 °C and the supernatant was discarded. The resin was then resuspended in 700 µL lysis buffer #2 and transferred to a Zymo-Spin P1 column (Zymo research). To remove non-specific contaminants, the resin was washed 3 times with 700 µL lysis buffer #2 prior to incubation with 100 µL elution buffer #2 (100 mM HEPES, pH 7.5, 1 M NaCl, 5 % glycerol, 500 mM imidazole) for 5 min on ice. Proteins were eluted by centrifugation 4,000 g for 1 min at 4 °C. Protein samples were analyzed on SDS-PAGE.

For preparative purifications of Strep-tagged gp44 and its variants, pRSF-Duet-derived plasmids encoding the *ncRNA*-*gp44* genes were transformed in *E. coli* BL21 Star (DE3) (Invitrogen) and plated on LB-agar plates containing kanamycin (Kan, 50 µg/mL). 10 mL LB-Kan overnight cultures (Kan, 50 µg/mL), grown shaking at 37 °C, were used to inoculate 2 L LB-Kan (Kan, 50 µg/mL) and incubated shaking at 200 r.p.m and 37 °C until the optical density at 600 nm (OD_600_) reached ≈ 0.6. Gene expression was induced with 1 mM IPTG, prior to incubation for 16 h at 16 °C. Cells were subsequently collected by centrifugation and resuspended in a lysis buffer #3 (100 mM HEPES, pH 7.5, 1 M NaCl, 5 % glycerol) prior to cell disruption using a microfluidizer. The lysate was clarified by centrifugation at 20,000 x g for 60 min at 4 °C. The supernatant was then loaded on a 1 mL StrepTrap XP column (Cytiva) pre-equilibrated in lysis buffer #3, subsequently washed with 20 CV lysis buffer #3 and eluted with 6 CV elution buffer #3 (100 mM HEPES, pH 7.5, 1 M NaCl, 5 % glycerol and 50 mM biotin). The sample was further purified by SEC using a Superose 75 10/300 (Cytiva) column pre-equilibrated in SEC buffer #2 (100 mM HEPES, pH 7.5, 1 M NaCl, 5 % glycerol). The protein was then concentrated to approximately 100 µL and the concentration was estimated based on the absorbance at 280 nm using a NanoDrop Eight spectrophotometer (Thermo Scientific). The sample was split in aliquots, snap frozen in liquid nitrogen and stored at −70 °C.

### Complex reconstitution

DRT4 WT or DRT4 ΔCTD were combined with gp44 WT or its D152G variant at a 1:4 ratio (5 µM DRT4 WT or DRT4 ΔCTD, 20 µM gp44 WT or gp44 D152G) in 1X rCutSmart buffer (NEB) to a final volume of 50 µL and incubated at 20°C for 10 min. For the complex reconstitution in presence of ssDNA production, 2 mM dNTPs (Thermo Fisher Scientific) were added to the same reaction mix and incubated at 37 °C for 10 min. Subsequently, reconstitution reactions or individual components were injected into a Separose 6 3.2/300, pre-equilibrated in a reconstitution buffer (10 mM Tris-HCl, pH 8.0, 50 mM NaCl). Peak fractions were analysed on SDS-PAGEs.

### *in vitro* ssDNA synthesis

0.52 µM DRT4 was combined with 250 µM dNTPs in 1X rCutSmart buffer (NEB) to a final volume of 30 µL and incubated at 37 °C. Reactions were stopped by adding 40 mM EDTA and incubated with 1 µL Proteinase K (∼20 mg/mL, Thermo Fisher Scientific) for 30 min at 37 °C. 10 µL of sample was mixed with 2 µL TriTrack DNA Loading Dye (Thermo Fisher Scientific) and analysed on 12 % polyacrylamide gels. Gels were stained with SYBR Gold (Thermo Fisher Scientific) and visualized using a GelDoc Go imaging system (Bio-Rad, software version 3.0.0.07).

### Oxford Nanopore sequencing of ssDNA and data analysis

DRT4 protein was purified as described above. 1.56 µM of DRT4 was combined with 2.5 mM of dNTPs in 1X rCutSmart buffer (NEB) to a final volume of 300 µL and incubated for 30 min at 37 °C. The reaction was stopped by adding 40 mM EDTA and subsequently incubated with 30 µL Proteinase K (∼20 mg/mL, Thermo Fisher Scientific) for 30 min at 37 °C. Nucleic acids were purified using a GeneJET PCR Purification Kit according to the manufacturer’s manual (Thermo Fisher Scientific). The concentration of nucleic acids was estimated based on the absorbance at 260 nm using a NanoDrop Eight spectrophotometer (Thermo Scientific). 650 ng of purified ssDNA was mixed with 0.5 mM dGTP (Thermo Fisher Scientific), 30 U TdT (Thermo Fisher Scientific) and 1X TdT reaction buffer (Thermo Fisher Scientific) in a final volume of 50 µL. The reaction was incubated for 45 min at 37 °C and stopped by adding 5 µL of 0.5 M EDTA and subsequently purified using a GeneJET PCR Purification Kit (Thermo Fisher Scientific). The purified sample was mixed with 0.5 µL of Phusion Plus DNA polymerase (Thermo Fisher Scientific), 0.2 mM dNTPs (Thermo Fisher Scientific), 1X Phusion Plus DNA polymerase buffer (Thermo Fisher Scientific) and 0.5 mM MJ508 primer in a final volume of 50 µL. The reaction was incubated for 30 s at 98 °C, 10 s at 60 °C, and 5 min at 72 °C in a ProFlex PCR system (Thermo Fisher Scientific) and subsequently purified with GeneJET PCR Purification Kit (Thermo Fisher Scientific).

The resulting dsDNA was used as input for Oxford Nanopore (ONT) sequencing library preparation. Ligation adapters (LA) were added using the Oxford Nanopore Technologies Ligation Sequencing Kit V14 (SQK-LSK114) according to manufacturer’s instructions (Manual GDE_9161_v114_revAB_26Aug2025). The sample was sequenced on a Flongle R10.4.1 flow cell (FLO-FLG114) and basecalled using the super accuracy model v5.2.0 on device (MinKNOW software version 25.09.16) without barcode and adapter trimming enabled.

Reads were first filtered to exclude reads without the Oxford Nanopore adapter sequence using seqkit grep with parameters “-p CCTGTACTTCGTTCAGTTACGTATTGCT -m 2” (v2.1.0; https://bioinf.shenwei.me/seqkit/note/), then the “SQK-LSK114” adapters were trimmed using dorado (v1.0.0; https://github.com/nanoporetech/dorado) and the longest 70% of reads were selected using filtlong (v0.2.1; https://github.com/rrwick/filtlong). The resulting reads were further inspected by eye and were AG-or CT-rich. CT rich reads (>70% CT) were changed to the reverse complement using a custom python script, because the ONT sequencing adapter ligates to both ends of DNA and both strands of the duplex can be sequenced. The reads were aligned using the MAFFT (v7.525; https://mafft.cbrc.jp/alignment/software/) algorithm and reads consisting of ligated random DNA, adapter fragments, or seemingly not fitting the alignment were removed using the seqkit tool. Since consecutive Gs were added by TdT at the 3’ end for dsDNA synthesis priming, 3’ ends with 6 or more consecutive Gs were trimmed using cutadapt ^41^(v5.2). The final MAFFT alignment consisted of 103 reads that were used to calculate nucleotide frequencies using a custom python script.

“Claude 3.5” Sonnet model was used to suggest python scripts and debug code, all generated code was manually inspected, edited and tested by a human. The final alignment was visualized using Ugene v51.0 software and saved in png format. The full analysis pipeline is available at https://data.mendeley.com/preview/yg4m99pwmn?a=3fb1b3fd-8b59-499f-80b1-a1d49e68bb38. Trimal (v1.5) was used to trim the alignment using the default preset “-automated1” and visualized by ugene software.

### ssDNA composition analysis by HPLC/ESI-MS

DRT4 sample was purified by the StrepTrap XP column (Cytiva) as described above. 1.5 mL of DRT4 (∼10 µM) was digested with a 100 µL of Proteinase K (∼20 mg/mL, Thermo Fisher Scientific) for 1 h at 37 °C. To purify the ssDNA, the resulting sample was loaded on a Superose 75 10/300 column (Cytiva), pre-equilibrated in a buffer containing 10 mM Tris-HCl pH 8.0 and 50 mM NaCl, for size-exclusion chromatography. The peak fraction was collected and the concentration was estimated based on the absorbance at 260 nm using a NanoDrop Eight spectrophotometer (Thermo Scientific). 100 pmol of nucleic acid sample was then combined with 2 µL of nucleoside digestion enzyme mix (NEB) and 1X reaction buffer (NEB) to a final volume of 40 µL and incubated for 1 h at 37 °C, followed by heat inactivation for 20 min at 80 °C.

Nucleotides were then analysed on an integrated HPLC/ESI-MS system (Agilent 1290 Infinity/ Q-TOF 6520) equipped with a Supelco Discovery HS C18 column (7.5 cm × 2.1 mm, 3 μm) by elution with a linear gradient of solvents A (0.0075% formic acid in water) and B (0.0075% formic acid in acetonitrile) at a flow of 0.3 mL/min at 30 °C as follows: 0–5 min, 0% B; 5–15 min, 10% B; 15–20 min, 98% B. Mass spectrometer was operating in the positive ion mode. High-resolution mass spectra of compounds were acquired in the 50–2000 m/z range. Ionization capillary voltage was set to 2500 V, fragmentor – to 150software version The results were analyzed with Agilent MassHunter Qualitative Analysis B.05.00 software.

### Cryo-EM grid preparation, data collection and data processing

DRT4 samples, and DRT-gp44 complexes, were purified as described above. For the nucleotide-bound state, 5 mM of dAPCPP were added to the purified sample and incubated for 10 min at room temperature before freezing grids.

Quantifoil R 1.2/1.3 300 copper mesh grids with a 2 nm continuous carbon support layer were glow-discharged at 20 mA for 7 s (GLOQUBE PLUS, Quorum) before applying 3 µL of sample (0.75 mg/mL). The grids were flash-frozen in liquid ethane using Vitrobot Mark IV (Thermo Scientific) with 100% relative humidity at 4 °C, with no preincubation and a blotting time of 5 s.

Datasets were collected using a Glacios Cryo-TEM (Thermo Scientific) operated at 200 kV and equipped with a Falcon III directed electron detector (Thermo Scientific) at ×92,000 magnification, with a pixel size of 1.1 Å, 30 frames with a total dose of 30 e−/Å2, and defocus range of −2.2 µm to −1.0 µm. Data collection was performed using EPU software version 3.2 and 3.13 (Thermo Fisher Scientific).

For the DRT4 apo sample, 2,086 images were recorded and patch motion correction and CTF estimation were performed on-the-fly using CryoSPARC Live (software version 4.2.1). Blob picking and particle extraction resulted in 1,708,489 particles which were sorted by two-dimensional (2D) classification. The initial model was generated and further processed using local and global CTF refinements, local motion correction and homogenous refinement, processing was performed using CryoSPARC ^42^ (software version 4.2.1). The final reconstruction was obtained with a global resolution of 2.23 Å containing 1,210,288 particles (D3 symmetry). Map sharpening was performed using Sharpening tools in CryoSPARC ^42^.

Data for DRT4 with bound dAPCPP were processed using CryoSPARC ^42^ (software version 5.0.2). A total of 2,368 images were recorded, patch motion correction and patch CTF estimation jobs and 2,041,746 particles were picked using blob picker and sorted by 2D classification. The initial models were generated and further classified using heterogeneous refinement. The best class was selected and further polished using local CTF refinement, reference based motion correction and iterative homogenous and non-uniform refinement jobs. The final reconstruction with a global resolution of 2.37 Å containing 1,162,671 particles (D3 symmetry) was obtained. Map sharpening was performed using Sharpening tools in CryoSPARC ^42^.

A total of 1,014 micrographs were collected for DRT4 in complex with gp44. Patch motion correction and CTF estimation were performed using CryoSPARC Live (software version 5.0.2). Blob picker was used for particle picker 518,546 particles were sorted by 2D classification. The initial models were generated and further classified using heterogenous refinement. Homogenous refinement of the best class was performed and the final reconstruction with a global resolution of 6.32 Å containing 104,808 particles was obtained.

### Model building, refinement, and figure preparation

The starting model of DRT4 was generated using AlphaFold2 ^43^ under the ColabFold ^43,44^ framework with default parameters and fitted into the Cryo-EM map in UCSF ChimeraX (software version 1.6.1). The model was modified and refined using Coot ^43–45^ (software version 0.9.8.7).

The model was then refined using Phenix real space refine job ^46^ with global minimization and local grid search strategies (software version 1.20.1–4487). The resulting model was used as a starting model for DRT4 bound to dAPCPP. The model was fitted into a map using UCSF ChimeraX (software version 1.10), further modified and refined in Coot (software version 0.9.8.1). Restrains for dAPCPP nucleotide were generated using AceDRG ^47^ (software version 330) as implemented in the CCP4 suite^48^ (software version 9.0) before fitting into the map in Coot. The model was refined using Phenix (software version 1.21.2-5419) as described above. Images were prepared using UCSF ChimeraX (software version 1.6.1).

### AlphaFold structure prediction

Models were predicted using AlphaFold 3 ^49^ (default settings) via the AlphaFold Server (https://deepmind.google/technologies/alphafold/alphafold-server/). pLDDT values were mapped on structures using UCSF ChimeraX (software version 1.6.1).

### Efficiency of transformation assay

DRT4-encoding plasmids were transformed in E. coli MG1655ΔRM and plated on LB-agar plates containing Cm (25 μg/mL) and incubated at 37 °C overnight. The following day single colonies were inoculated into 4 mL LB medium and grown shaking at 37 °C to prepare starter cultures for electrocompetent cell preparation.

The overnight culture was then used to inoculate 50 mL of LB medium, which was incubated at 37 °C with shaking until the culture reached an OD600 of 0.35. The culture was immediately transferred to ice and chilled for 30 min. All subsequent steps were performed at 4°C using pre-chilled solutions and centrifuge tubes. Cells were harvested by centrifugation at 1,000 × g for 20 min at 4 °C. The supernatant was discarded, and the cell pellet was gently resuspended in 10 mL of ice-cold sterile H_2_O. Cells were collected again by centrifugation at 1,000 × g for 20 min at 4 °C. The supernatant was removed, and the pellet was resuspended in 5 mL of ice-cold sterile H_2_O, followed by a second centrifugation at 1,000 × g for 20 min at 4 °C. The resulting pellet was gently resuspended in 2 mL of ice-cold 10% glycerol and transferred to a pre-chilled microcentrifuge tube. Cells were harvested by centrifugation at 1,000 × g for 20 min at 4 °C. The supernatant was carefully removed, and the pellet was gently resuspended in 50 µL of ice-cold 10% glycerol. The resulting electrocompetent cells were aliquoted and snap-frozen in liquid nitrogen before storage at −80°C.

Aliquoted electrocompetent cells (50 µL) were thawed on ice, and 10 ng of the *ncRNA-gp44* plasmid was added. The cell-DNA suspension was then transferred into a pre-chilled 1 mm electroporation cuvette (Fisher scientific) and electroporated using a MicroPulser (Bio-Rad). Immediately thereafter, cells were recovered in 1 mL of LB shaking at 37 °C for 1 h. Cells were plated on LB agar supplemented with Cm (25 μg/mL) and Amp (50 μg/mL) and incubated overnight at 37 °C. To induce gp44 expression, the plates additionally contained 0.2 % arabinose (Carl Roth). Transformation efficiency was calculated with the formula: (number of colonies × total recovery volume / volume plated) / amount of DNA (µg). Data were analyzed and plotted in Prism (software version 10.6.1 (799), Graphpad).

### RNase activity assay

Reactions were assembled by combining 32 nM DRT4 WT or DRT4 ΔCTD with 128 nM gp44 and 100 nM U(5)GU(5) RNA FQ probe (IDT, Supplementary table 2) and 1X rCutSmart reaction buffer (NEB) in a final volume of 30 µL in 384-well flat bottom black polystyrene assay plates (Thermo Fisher Scientific). For ssDNA synthesis, either 10.4 µM dATP (Thermo Fisher Scientific), dGTP (Thermo Fisher Scientific), dAPCPP (Jena bioscience), dGPCPP (Jena bioscience), or a mix of dNTPs (Thermo Fisher Scientific) were used. The reactions were monitored by fluorescence measurements (*λ*_ex_: 489 nm; *λ*_em_: 522 nm) in a CLARIOstar plus plate reader (BMG Labtech, software version 5.70 R2) at 37 °C every 30 sec. Data were analyzed and plotted in Prism (software version 10.6.1 (799), Graphpad).

### Protein analyses by SDS-PAGE

12 µL of samples were mixed with 4 µL of 4x SDS dye (10% glycerol [v/v], 2% SDS [w/v], 63 mM Tris-HCl pH 6.8, 0.1% 2-Mercaptoethanol, 0.01% bromophenol blue [w/v]) and denatured for 5 min at 95 °C. Samples were separated in homemade 12% Tris-Glycine SDS polyacrylamide gels and stained with Coomassie solution (0.1% Coomassie R-250, 40% [v/v] EtOH, 10% [v/v] acetic acid). Gels were visualized using a GelDoc Go imaging system (Bio-Rad, software software version 3.0.0.07).

## Data Availability

Cryo-EM maps and models, as well as other data are available from the authors upon reasonable request and with the final version of the peer reviewed and published manuscript.

## Acknowledgements

We thank the department of Protein - DNA interactions at the Life Sciences Center at Vilnius University (Virginijus Šikšnys) for access to equipment and the Vilnius University cryo-EM facility (Giedrius Sasnauskas) for support. The authors thank Giedrė Tamulaitienė for critical reading and comments on the manuscript. We acknowledge Audronė Rukšėnaitė (Vilnius University, Lithuania) for technical support. The authors thank Alexander Harms (ETH Zürich, Switzerland) for providing the BASEL phage collection, and Inga Songalienė (Vilnius University, Lithuania) for discussion. We acknowledge Linyi Gao and Feng Zhang (MIT, USA) for kindly providing pLG012 (Addgene plasmid #157890). P.P. receives funding from the EMBC under EMBO Installation Grant agreement number 5342-2023 (P.P.), and the RCL under Measure No. 12-001-01-01-01, Project No: S-A-UEI-23-10 (P.P.). J.J. was supported by Vilnius University Science Promotion fund project MSFJM-12/2021. The funders had no role in study design, data collection and analysis, decision to publish or preparation of the manuscript.

## Author Contributions

P.P. conceived the study. M.J. performed growth and infection assays, produced and purified proteins, performed ssDNA production, RNA cleavage assays and complex reconstitution *in vitro*. A. C. and K.K. tested protection against BASEL collection and isolated phage escapers with guidance by J.J. and M.J.. K.K. and M.J. cloned constructs. M.J., A.S. and A.M. prepared cryo-EM samples. A.S. and A.M. collected and processed cryo-EM data with input from P.P. and L.M.. K.K, A.M and A.S. built and refined structure models. J.J. sequenced phage escaper genomes and ssDNA. A.C. performed homolog search of gp44 with guidance by J.J., and performed EOP experiments testing DRT4 variants for antiphage defense with guidance by M.J.. M.T. performed the HPLC/ESI-MS analysis. All authors designed experiments and analyzed data. M.J. and P.P. wrote the manuscript and prepared figures with input from all authors. The manuscript was reviewed and approved by all authors.

## References

1. Hampton, H. G., Watson, B. N. J. & Fineran, P. C. The arms race between bacteria and their phage foes. Nature 577, 327–336 (2020).

2. Lopatina, A., Tal, N. & Sorek, R. Abortive Infection: Bacterial Suicide as an Antiviral Immune Strategy. Annu Rev Virol 7, 371–384 (2020).

3. Labrie, S. J., Samson, J. E. & Moineau, S. Bacteriophage resistance mechanisms. Nat Rev Microbiol 8, 317–327 (2010).

4. Menéndez-Arias, L., Sebastián-Martín, A. & Álvarez, M. Viral reverse transcriptases. Virus Res 234, 153– 176 (2017).

5. Baltimore, D. RNA-dependent DNA polymerase in virions of RNA tumour viruses. Nature 226, 1209– 1211 (1970).

6. González-Delgado, A., Mestre, M. R., Martínez-Abarca, F. & Toro, N. Prokaryotic reverse transcriptases: from retroelements to specialized defense systems. FEMS Microbiol Rev 45, (2021).

7. Mestre, M. R. et al. UG/Abi: a highly diverse family of prokaryotic reverse transcriptases associated with defense functions. Nucleic Acids Res 50, 6084–6101 (2022).

8. Figiel, M. et al. Mechanism of protein-primed template-independent DNA synthesis by Abi polymerases. Nucleic Acids Res 50, 10026–10040 (2022).

9. Gapińska, M. et al. Structure-functional characterization of Lactococcus AbiA phage defense system. Nucleic Acids Res 52, 4723–4738 (2024).

10. Wang, Y., Wu, H., Li, J., Tian, Z. & Deng, Z. Cryo-EM structure of the bacterial anti-phage defense system DRT6. Biochem Biophys Res Commun 791, 152955 (2025).

11. Wang, Y. et al. Anti-phage defense mechanism involving phage-encoded DNA binding protein and bacterial reverse transcriptase DRT4. Nat Commun 17, 289 (2026).

12. Rong, X. et al. DNA polymerization activates RNA cleavage of a reverse transcriptase-like antiviral enzyme. Science 393, eaef3178 (2026).

13. Figiel, M. et al. Structures and enzymatic mechanisms of DRT7/UG10 antiphage reverse transcriptases. bioRxiv (2026) doi:10.64898/2026.02.16.706125.

14. Stokar-Avihail, A. et al. Discovery of phage determinants that confer sensitivity to bacterial immune systems. Cell 186, 1863–1876.e16 (2023).

15. Gao, L. et al. Diverse enzymatic activities mediate antiviral immunity in prokaryotes. Science 369, 1077– 1084 (2020).

16. Humolli, D. et al. Completing the BASEL phage collection to unlock hidden diversity for systematic exploration of phage-host interactions. PLoS Biol 23, e3003063 (2025).

17. Weinberg, Z. et al. Detection of 224 candidate structured RNAs by comparative analysis of specific subsets of intergenic regions. Nucleic Acids Res 45, 10811–10823 (2017).

18. Kazlauskas, D. & Venclovas, C. Two distinct SSB protein families in nucleo-cytoplasmic large DNA viruses. Bioinformatics 28, 3186–3190 (2012).

19. Arcus, V. OB-fold domains: a snapshot of the evolution of sequence, structure and function. Curr Opin Struct Biol 12, 794–801 (2002).

20. Cernooka, E., Rumnieks, J., Tars, K. & Kazaks, A. Structural Basis for DNA Recognition of a Single-stranded DNA-binding Protein from Enterobacter Phage Enc34. Scientific Reports 7, 15529 (2017).

21. Pata, J. D., Yin, Y. W. & Lahiri, I. Editorial: Nucleic Acid Polymerases: The Two-Metal-Ion Mechanism and Beyond. Front Mol Biosci 9, 948326 (2022).

22. Helmann, J. D. Metals in Motion: Understanding Labile Metal Pools in Bacteria. Biochemistry 64, 329–345 (2025).

23. Gong, S. et al. Kinetic and thermodynamic analysis defines roles for two metal ions in DNA polymerase specificity and catalysis. J Biol Chem 296, 100184 (2021).

24. Huiting, E. & Bondy-Denomy, J. Defining the expanding mechanisms of phage-mediated activation of bacterial immunity. Curr Opin Microbiol 74, 102325 (2023).

25. Aframian, N. & Eldar, A. Abortive infection antiphage defense systems: separating mechanism and phenotype. Trends Microbiol 31, 1003–1012 (2023).

26. Vashishtha, A. K., Wang, J. & Konigsberg, W. H. Different Divalent Cations Alter the Kinetics and Fidelity of DNA Polymerases. J Biol Chem 291, 20869–20875 (2016).

27. Vaisman, A., Ling, H., Woodgate, R. & Yang, W. Fidelity of Dpo4: effect of metal ions, nucleotide selection and pyrophosphorolysis. EMBO J 24, 2957–2967 (2005).

28. Yu, C. et al. Phage SSB detection by retron Eco8 msDNA unleashes nuclease-mediated immunity. Mol Cell 85, 4243–4253.e4 (2025).

29. Ji, C.-G. et al. Mechanistic insights into activation of bacterial Retron-Eco8 immunity by phage protein SSB. Nat Commun 17, (2026).

30. Xiong, C. et al. Structural basis of Retron-Eco8-mediated antiphage defense. Nucleic Acids Res 54, (2026).

31. Hooper, M. M. et al. Phage-encoded factor stimulates DNA degradation by the Hna anti-phage defense system. Nat Commun 17, (2026).

32. Bari, S. M. N. et al. A unique mode of nucleic acid immunity performed by a multifunctional bacterial enzyme. Cell Host Microbe 30, 570–582.e7 (2022).

33. Tuck, O. T. et al. Genome integrity sensing by the broad-spectrum Hachiman antiphage defense complex. Cell 187, 6914–6928.e20 (2024).

34. Tang, Y., Liu, T., Xiong, C., Chen, Q. & Yu, Y. Oligomer disassembly activates an HEPN-containing bacterial defense system. Mol Cell 86, 2560–2571.e5 (2026).

35. Puteikiene, R., et al. Viral SSB-bound ssDNA activates the bacterial anti-phage defense system DARNA. bioRxiv (2026) doi:10.64898/2026.02.22.705581.

36. Emond, E. et al. Phenotypic and genetic characterization of the bacteriophage abortive infection mechanism AbiK from Lactococcus lactis. Appl Environ Microbiol 63, 1274–1283 (1997).

37. Maffei, E. et al. Systematic exploration of Escherichia coli phage-host interactions with the BASEL phage collection. PLoS Biol 19, e3001424 (2021).

38. Jasnauskaitė, M. et al. Structure and mechanism of antiphage retron Eco2. Nat Struct Mol Biol 33, 330–340 (2026).

39. Ewels, P., Magnusson, M., Lundin, S. & Käller, M. MultiQC: summarize analysis results for multiple tools and samples in a single report. Bioinformatics 32, 3047–3048 (2016).

40. Deatherage, D. E., Traverse, C. C., Wolf, L. N. & Barrick, J. E. Detecting rare structural variation in evolving microbial populations from new sequence junctions using breseq. Front Genet 5, 468 (2014).

41. Martin, M. Cutadapt removes adapter sequences from high-throughput sequencing reads. EMBnet J. 17, 10 (2011).

42. Punjani, A., Rubinstein, J. L., Fleet, D. J. & Brubaker, M. A. cryoSPARC: algorithms for rapid unsupervised cryo-EM structure determination. Nat Methods 14, 290–296 (2017).

43. Cabrera, R. et al. Novel De Novo DLL4 Missense and Highly Accurate Protein Structure Prediction in Adams-Oliver Type 6 Syndrome. Clin Case Rep 13, e70933 (2025).

44. Mirdita, M. et al. ColabFold: making protein folding accessible to all. Nat Methods 19, 679–682 (2022).

45. Emsley, P., Lohkamp, B., Scott, W. G. & Cowtan, K. Features and development of Coot. Acta Crystallogr D Biol Crystallogr 66, 486–501 (2010).

46. Liebschner, D. et al. Macromolecular structure determination using X-rays, neutrons and electrons: recent developments in Phenix. Acta Crystallogr D Struct Biol 75, 861–877 (2019).

47. Long, F. et al. AceDRG: a stereochemical description generator for ligands. Acta Crystallogr D Struct Biol 73, 112–122 (2017).

48. Agirre, J. et al. The CCP4 suite: integrative software for macromolecular crystallography. Acta Crystallogr D Struct Biol 79, 449–461 (2023).

49. Abramson, J. et al. Accurate structure prediction of biomolecular interactions with AlphaFold 3. Nature 630, 493–500 (2024).

