## Supplementary Information for "Dual recognition of phage SSB triggers DRT4 for antiphage defense"

#### Table of Content

Supplementary Figures 1-15 pages 2-18

Supplementary Tables 1-4 pages 19-24

Supplementary References page 24

### Supplementary Figures

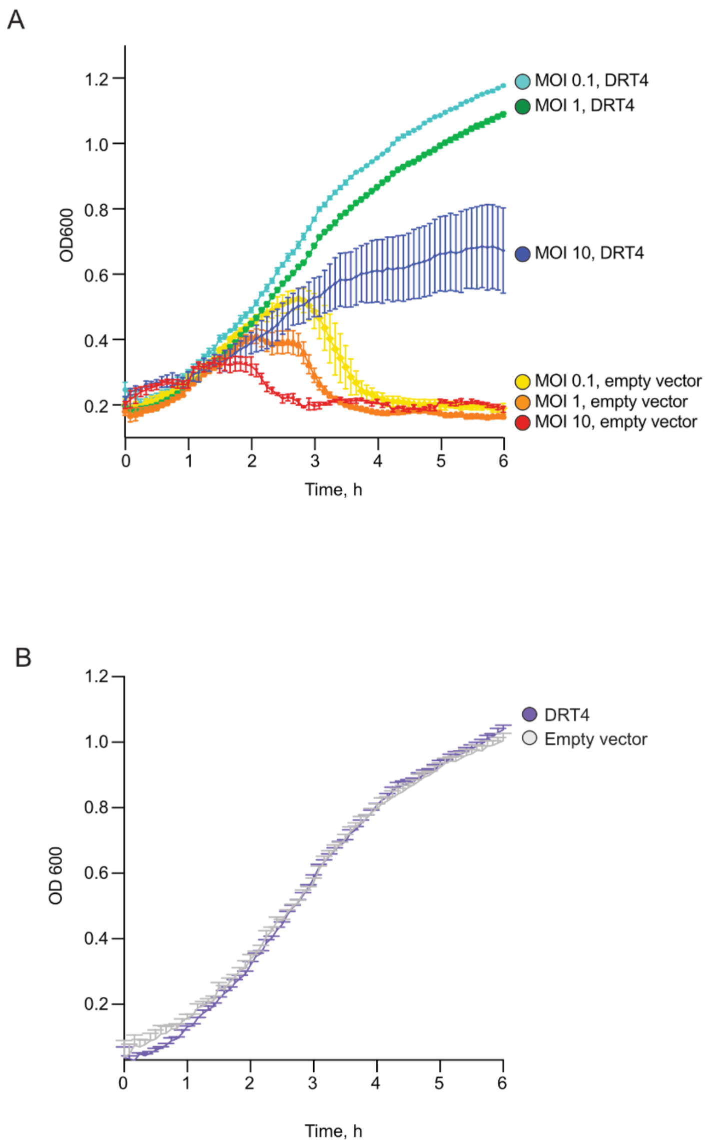


**Supplementary Fig. 1| DRT4 protects against phage T5 without impairing growth in the absence of infection. A,** Growth curves of *E. coli* expressing DRT4 or carrying an empty-vector control following T5 infection at the indicated multiplicities of infection (MOIs). *n* = 3 technical replicates; mean ± s.d.. **B,** Growth curves of the same strains in the absence of phage infection. *n* = 3 technical replicates; mean ± s.d..

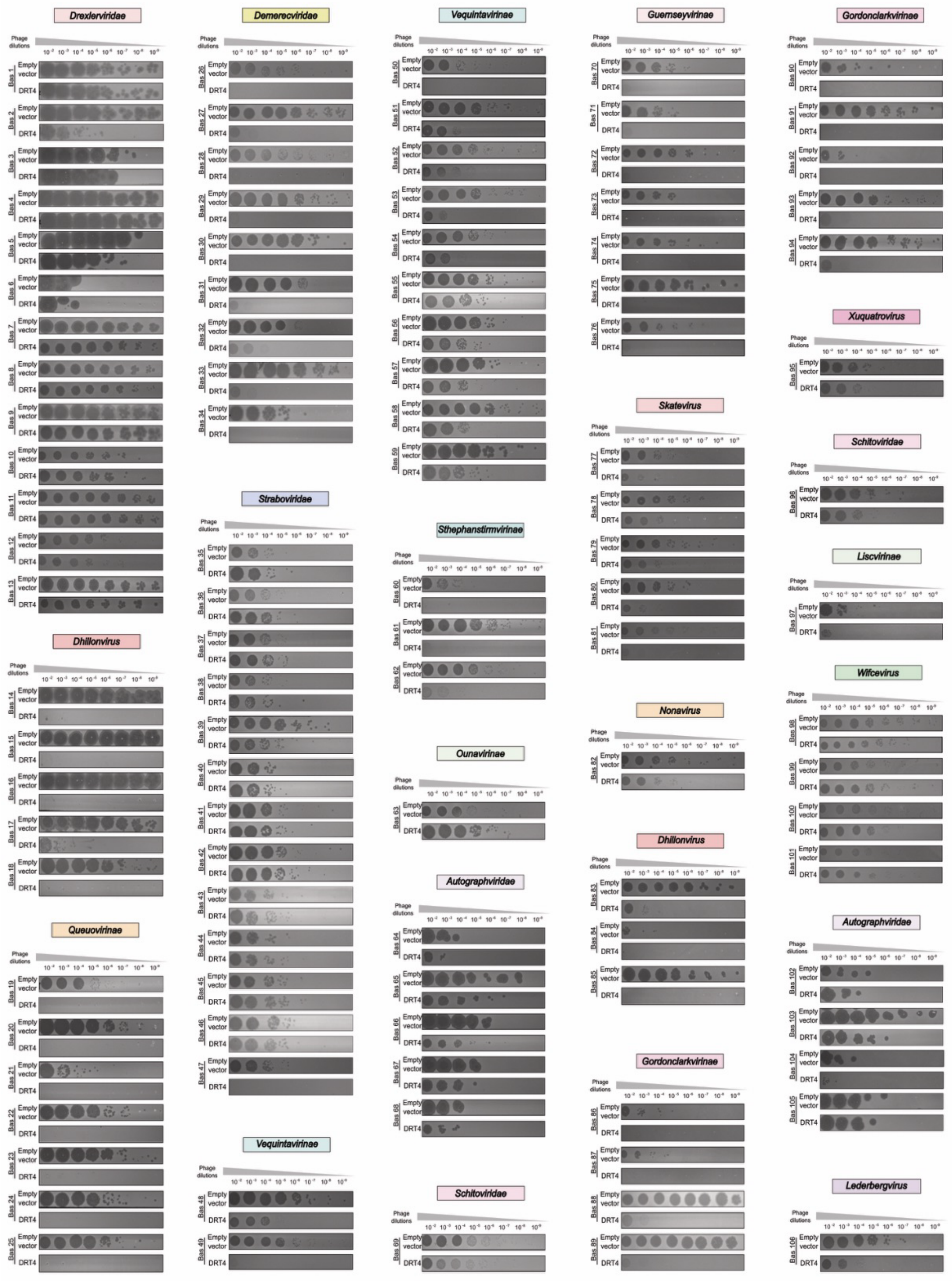


**Supplementary Fig. 2 | DRT4-mediated protection against phages from the BASEL collection.** Representative spot-plating assays of BASEL phages on *E. coli* carrying either the empty-vector control (pACYC184) or the DRT4 expression plasmid (pLG012 [1]). Representative data, $n = 3$ technical replicates.

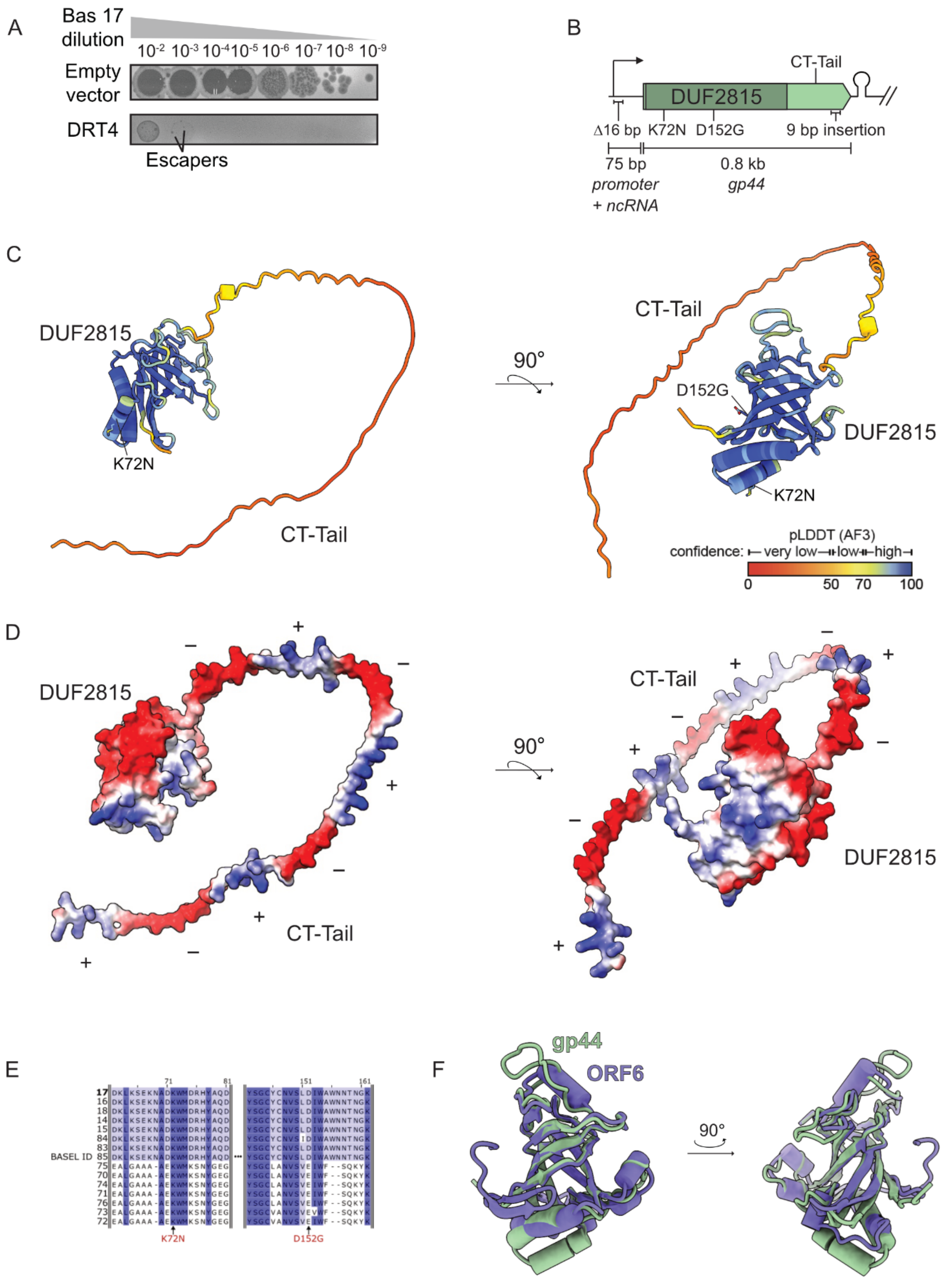


**Supplementary Fig. 3 | Bas17 escapes DRT4-mediated defense through mutations in the gp44 locus. A,** Representative spot-plating assays of Bas17 on *E. coli* carrying either the empty-vector control (pACYC184) or the DRT4 expression plasmid (pLG012; [1]). Escaper plaques are highlighted by lines. Representative data; $n \geq 2$ technical replicates. **B,** Schematic of the *gp44* locus showing escaper-associated mutations. **C,** AlphaFold 3 model of gp44, with residues substituted in escaper variants marked by lines. Below, the model colored by pLDDT score to indicate prediction confidence. **D,** Predicted gp44 structure shown as a molecular surface colored by electrostatic potential. Plus and minus signs indicate tandem-repeat segments enriched in positively charged residues (lysines and arginines) or negatively charged residues (aspartates and glutamates), respectively. **E,** Multiple sequence alignment of Bas17 gp44 and homologs encoded by other phages in the BASEL collection, shaded blue according to sequence identity. Arrows indicate the positions of the K72N and D152G escape substitutions (red labels). **F,** Structural superposition of ORF6 (OB fold, purple; PDB 5ODK; [2]) and the predicted gp44 DUF2815 domain (green). The root-mean-square deviation (RMSD) is 0.84 Å over 82 atom pairs retained after pruning and 8.5 Å over all 149 aligned atom pairs.

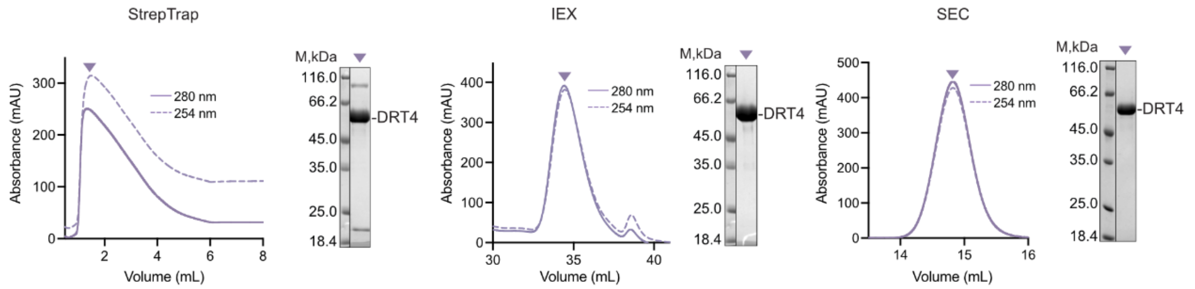


**Supplementary Fig. 4 | Purification of DRT4.** From left to right, chromatograms from StrepTrap affinity chromatography, ion-exchange chromatography (IEX), and size-exclusion chromatography (SEC) using a Superose 6 10/300 column, with corresponding SDS-PAGE analyses of the main peak fractions indicated by triangles. Representative data; $n \geq 2$ biological replicates.

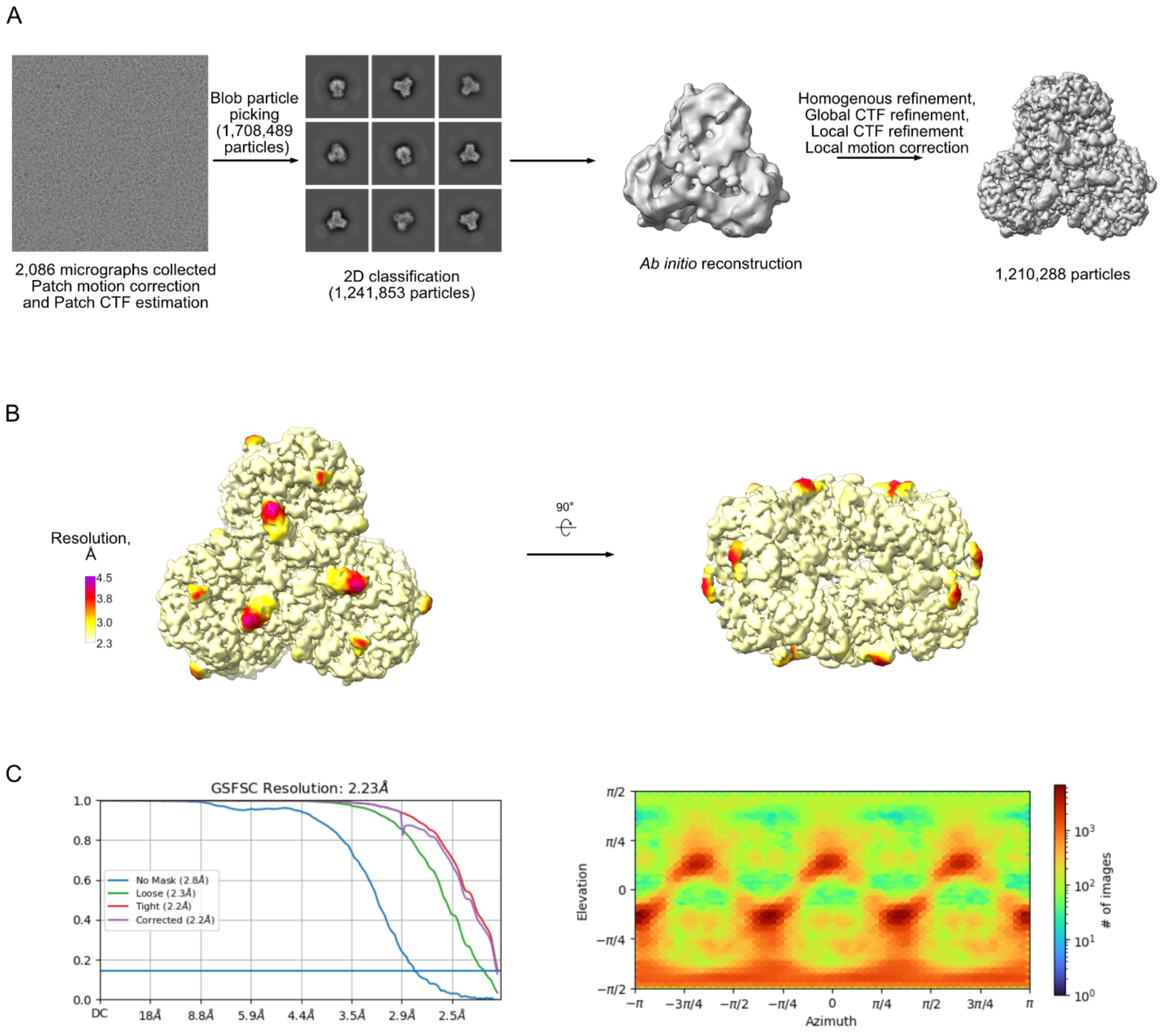


**Supplementary Fig. 5 | Cryo-EM data processing and reconstruction of DRT4. A,** Data-processing workflow in cryoSPARC. **B,** Cryo-EM map colored by local resolution, shown in two orientations. **C,** Fourier shell correlation (FSC) curves (left) and particle angular distribution plots (right), calculated in cryoSPARC.

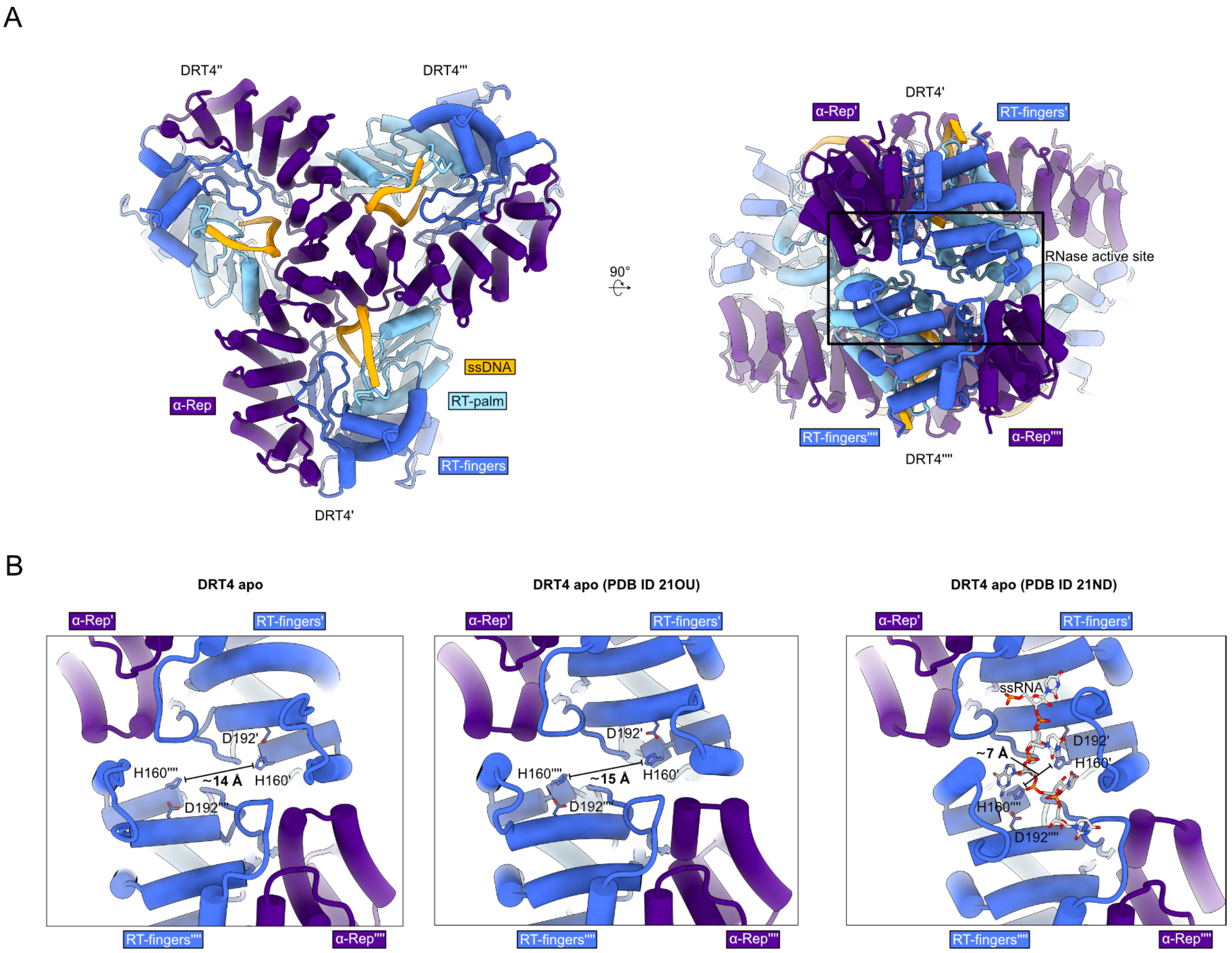


**Supplementary Fig. 6 | DRT4 adopts an RNase-inactive conformation. A,** Cryo-EM structure of DRT4 determined in this study, shown in 90°-rotated views. The box marks the RNase active site. **B,** Comparison of RNase active-site configurations in DRT4 from this study (left), a previously reported RNase-inactive DRT4 structure (PDB 21OU; [3]), and an RNase-active DRT4 structure bound to ssRNA (grey sticks) (PDB 24ND; [3]). Distances between histidine 160 residues in the juxtaposed RT fingers subdomains are indicated.

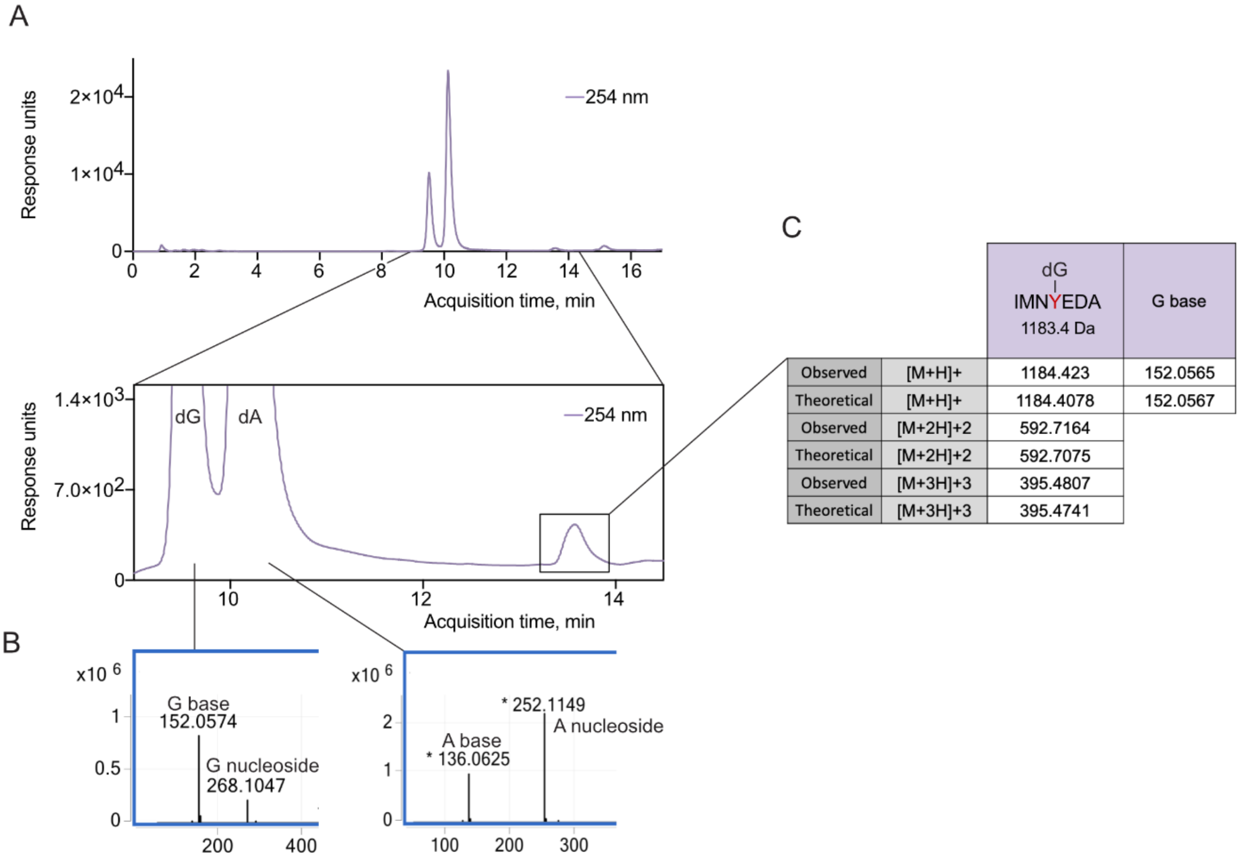


| | | dG–IMNYEDA 1183.4 Da | G base |
|---|---|---|---|
| Observed | [M+H]+ | 1184.423 | 152.0565 |
| Theoretical | [M+H]+ | 1184.4078 | 152.0567 |
| Observed | [M+2H]+2 | 592.7164 | |
| Theoretical | [M+2H]+2 | 592.7075 | |
| Observed | [M+3H]+3 | 395.4807 | |
| Theoretical | [M+3H]+3 | 395.4741 | |

**Supplementary Fig. 7 | Y125 is covalently linked to a dG residue. A,** Top: HPLC-MS chromatogram of products obtained after DRT4-oligo digestion with Proteinase K and nucleoside digestion enzyme mix. Bottom: a close-up of the two major peaks, dA and dG. The third minor peak, with a later retention time, is highlighted with a square. **B,** Extracted mass spectra at the two major peaks, dG and dA. **C,** Extensive mass spectrometry analysis of the minor peak. The peak represents a peptide: a seven amino acid sequence including the Y125, with a deoxyguanosine attached. The presence of G base was confirmed by the detection of its mass.

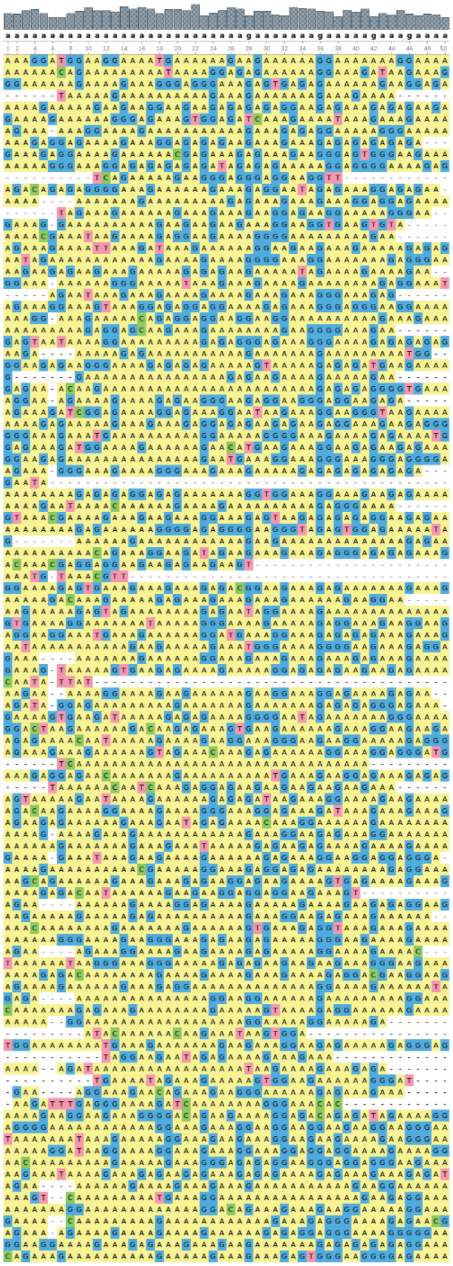

**Supplementary Fig. 8 | Alignment of ssDNA sequencing reads.** Gray bars above the alignment indicate conservation at each position, with the consensus sequence shown in black below the bars. Numbers indicate alignment positions. Nucleotides are colored as follows: dA, yellow; dG, blue; dC, green; and dT, red. $n = 1$.

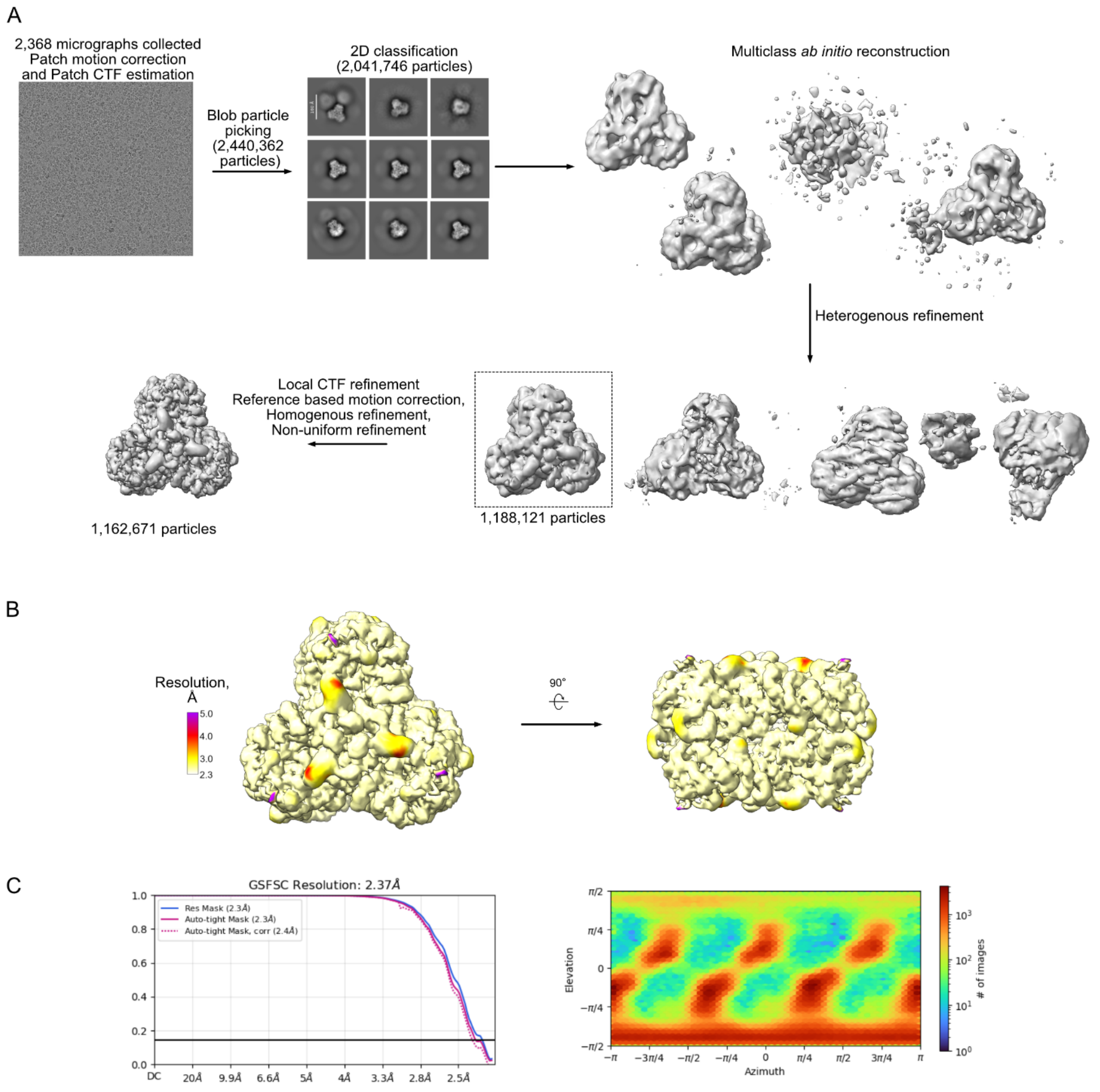


**Supplementary Fig. 9 | Cryo-EM data processing and reconstruction of dAPCPP-bound DRT4. A,** Data-processing workflow in cryoSPARC. **B,** Cryo-EM map colored by local resolution, shown in two orientations. **C,** Fourier shell correlation (FSC) curves (left) and particle angular distribution plots (right), calculated in cryoSPARC.

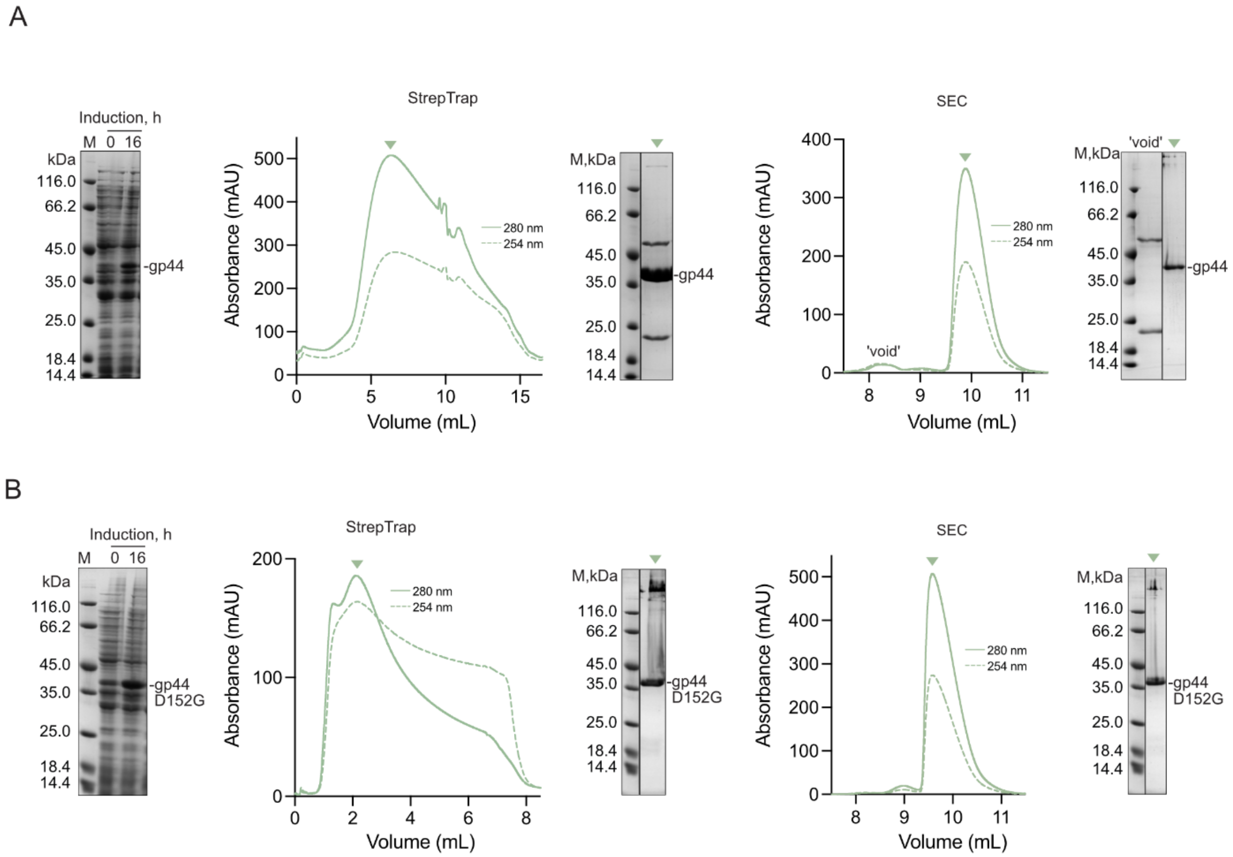


**Supplementary Fig. 10 | Expression and purification of gp44 and its D152G variant. A and B,** Expression and purification of wild-type gp44 (A) and gp44 D152G (B). Left, SDS-PAGE analysis of samples collected before and after induction. Middle, Strep-tag affinity chromatography traces and corresponding SDS-PAGE analyses of peak fractions indicated by triangles. Right, size-exclusion chromatography (SEC) traces obtained using a Superose 75 10/300 column and corresponding SDS-PAGE analyses of peak fractions indicated by triangles. Representative data; $n \geq 2$ biological replicates.

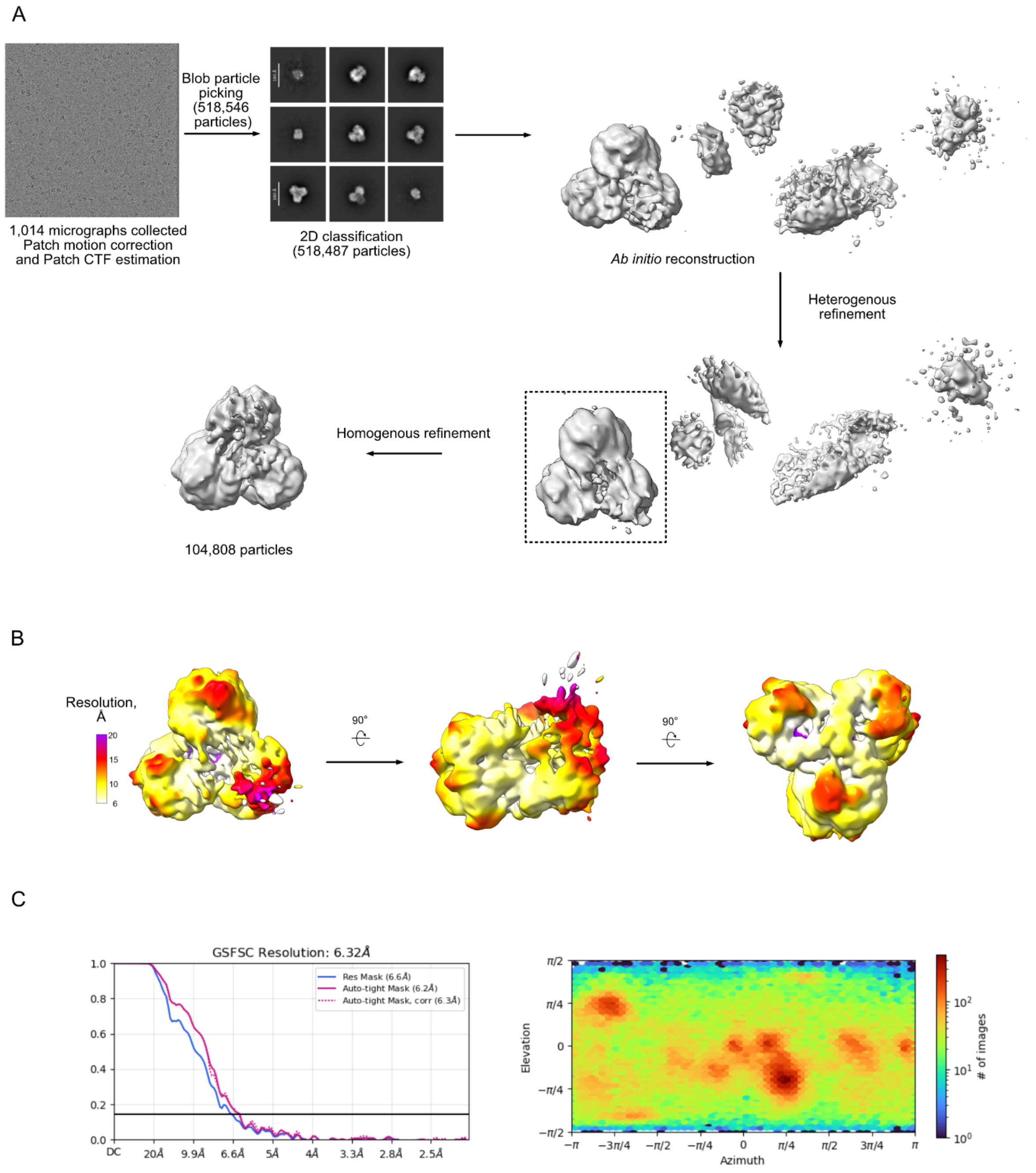


**Supplementary Fig. 11 | Cryo-EM data processing and reconstruction of DRT4 in the presence of gp44. A,** Data-processing workflow in cryoSPARC. **B,** Cryo-EM map colored by local resolution, shown in two orientations. **C,** Fourier shell correlation (FSC) curves (left) and particle angular distribution plots (right), calculated in cryoSPARC.

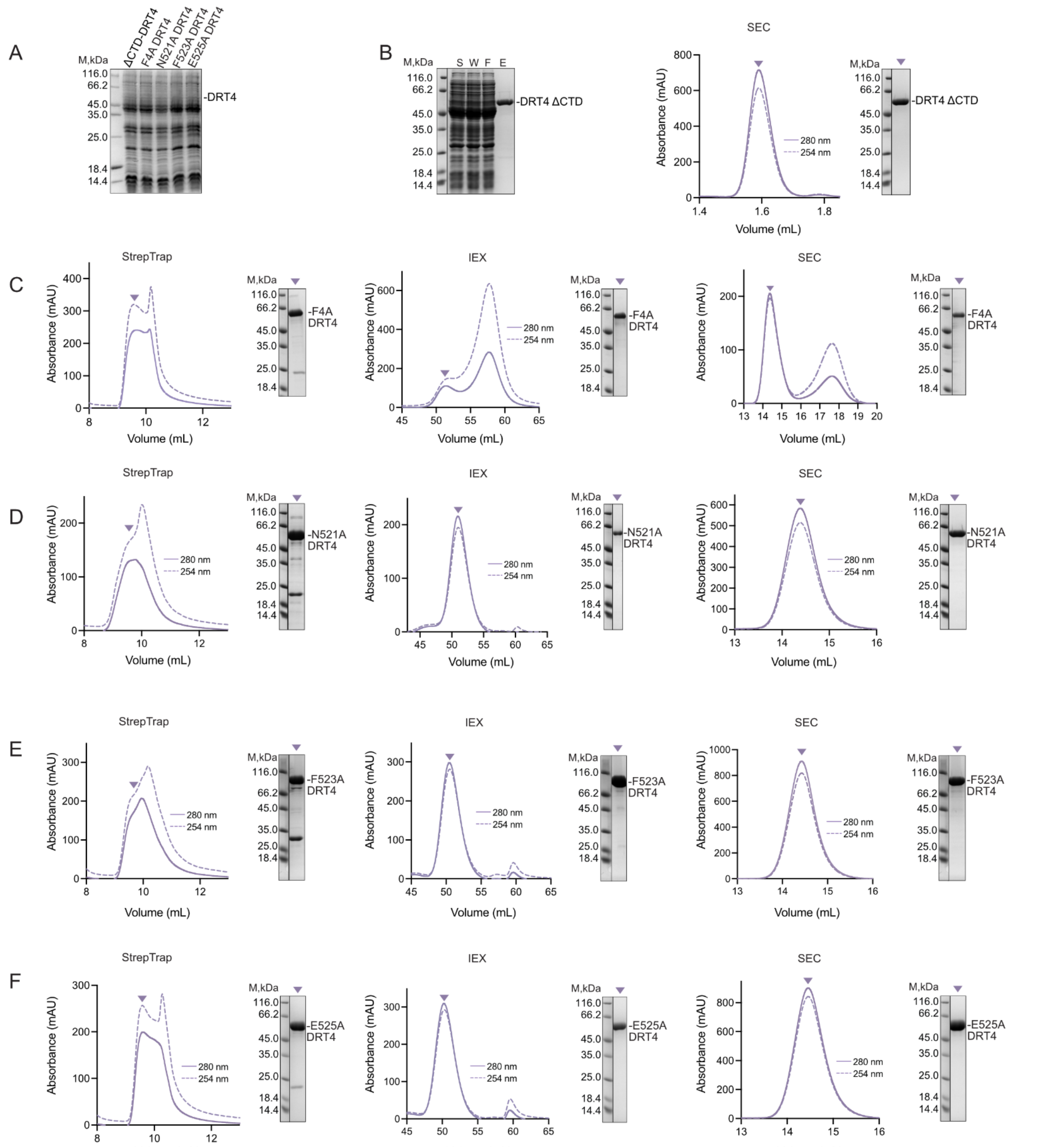


**Supplementary Fig. 12 | Expression and purification of DRT4 variants. A,** SDS-PAGE analysis of whole-cell protein samples following overnight incubation. **B,** Purification of DRT4 ΔCTD. Left, SDS-PAGE analysis of the soluble fraction (S) and wash (W), flow-through (F), and elution (E) fractions from Strep-tag affinity chromatography. Right, size-exclusion chromatography (SEC) trace obtained using a Superose 6 3.2/300 column and corresponding SDS-PAGE analysis. The triangle indicates the peak fraction. **C–F,** Purification of DRT4 variants F4A

(C), N521A (D), F523A (E), and E525A (F). From left to right, Strep-tag affinity chromatography, ion-exchange chromatography (IEX), and SEC traces, with corresponding SDS-PAGE analyses. SEC was performed using a Superose 6 10/300 column. Representative data; $n \geq 2$ biological replicates.

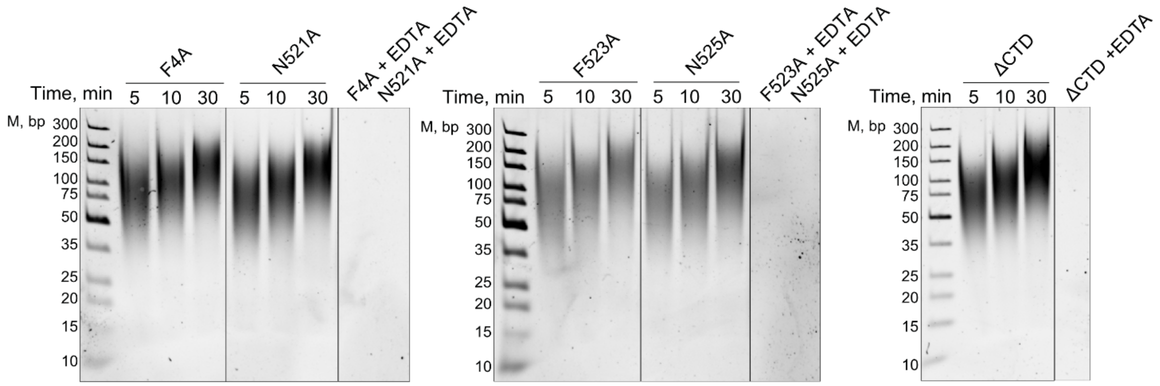


**Supplementary Fig. 13 | In vitro ssDNA synthesis by DRT4 variants.** Polyacrylamide gel analysis of ssDNA products synthesized by DRT4 variants at the indicated time points. Representative data; $n \geq 2$ biological replicates.

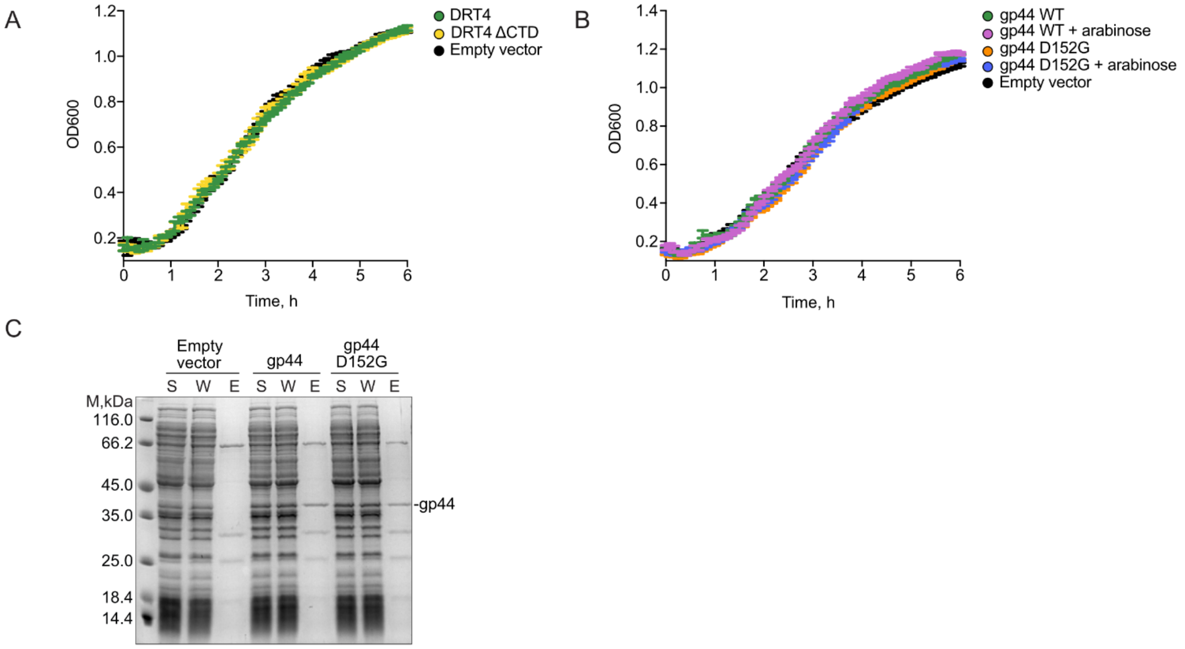


**Supplementary Fig. 14 | Effects of DRT4 variants and gp44 expression on bacterial growth, and purification of gp44. A,** Growth curves of *E. coli* carrying the empty-vector control (pACYC184), a DRT4 expression plasmid (pMJ016), or a DRT4 ΔCTD expression plasmid (pMJ018). **B,** Growth curves of *E. coli* carrying the empty-vector control (pBAD) or constructs encoding ncRNA-gp44 (pMJ024) or its D152G escape variant (pMJ025), with or without arabinose induction. Representative data, $n = 3$ technical replicates, mean ± s.d. **C,** SDS-PAGE analysis of soluble (S), wash (W), and elution (E) fractions from Ni-NTA affinity purification of gp44. Representative data; $n \geq 2$ biological replicates.

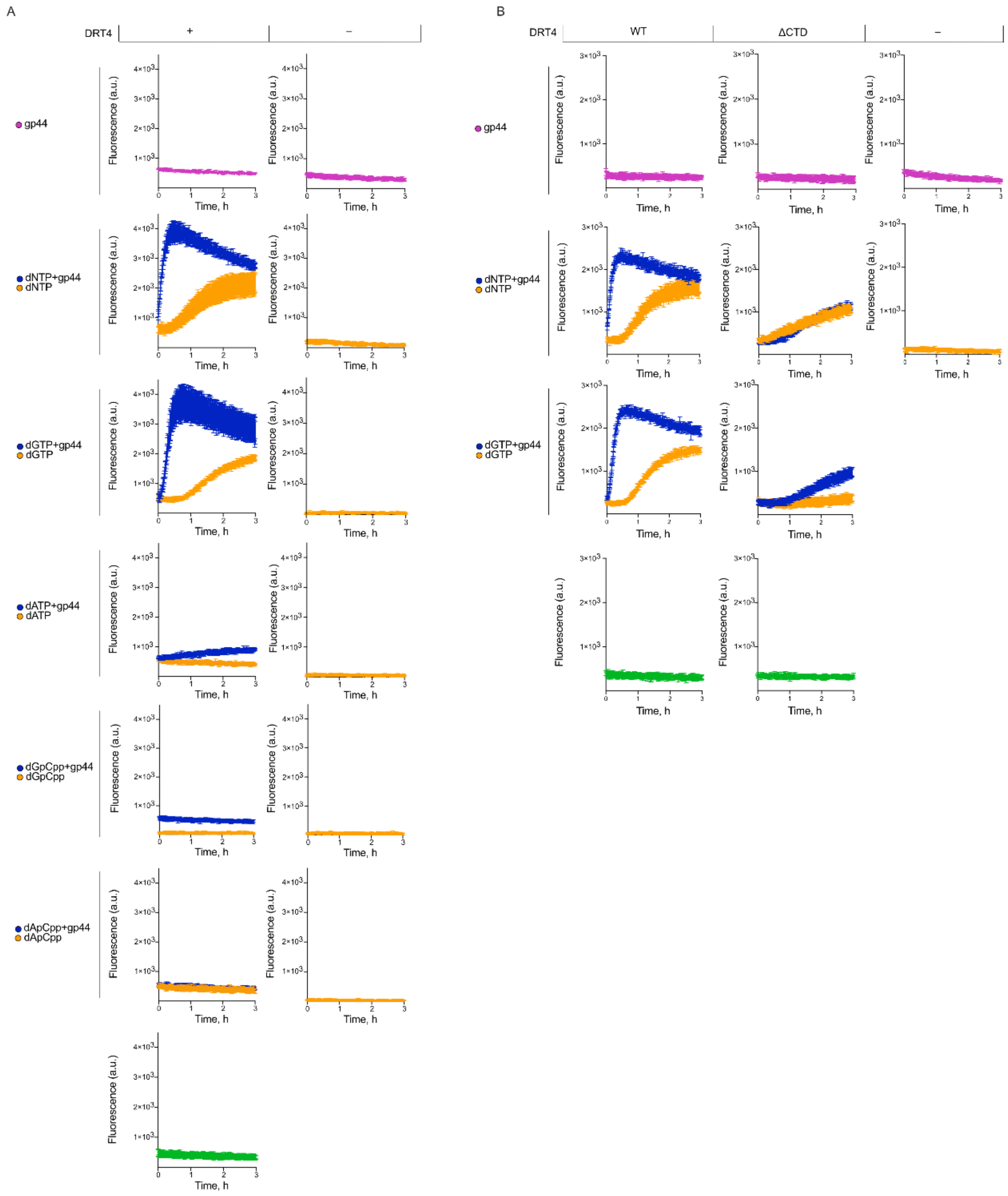


**Supplementary Fig. 15| Fluorophore–quencher (FQ) reporter assays of RNase activity. A,** Fluorescence traces for the indicated combinations of DRT4, gp44, and nucleotide substrates. technical replicates; mean ± s.d. **B,** Fluorescence traces for wild-type DRT4 or DRT4 ΔCTD in the presence of gp44 and either dGTP alone or all four dNTPs. $n = 3$ technical replicates; mean ± s.d.

### Supplementary Tables

**Supplementary Tab. 1: Plasmids**

| ID | Purpose | Features | Selection marker |
|---|---|---|---|
| pLG012 [1] (Addgene #157890) | Phage plaque assays | pACYC-derived vector encoding the DRT4 locus. | Chloramphenicol |
| pACYC184 (NEB) | Phage plaque assays; empty vector control | p15A-ori, cloning vector | Chloramphenicol |
| pBAD (Department of Protein-DNA interactions, Life Science Centre, Vilnius University) | Activity assays *in vitro*, protein purification; empty vector control | Cloning vector | Ampicillin |
| pMJ016 | Protein purification, activity assays *in vitro* and *in vivo,* phage plaque assays | Derivative of pLG012. Contains a Strep-tag II encoding sequence fused downstream to the *drt4* gene | Chloramphenicol |
| pMJ017 | Protein purification, activity assays *in vitro* and *in vivo,* phage plaque assays | Derivative of pLG012. Contains a Strep-tag II-encoding sequence fused downstream to the *drt4* gene (D240A, D241A mutant, dRT) | Chloramphenicol |
| pMJ018 | Protein purification, activity assays *in vitro* and *in vivo,* phage plaque assays | Derivative of pLG012. Contains a Strep-tag II-encoding sequence fused downstream to the *drt4* gene (mutant with deletion of 508-540 residues, ΔCTD-DRT4) | Chloramphenicol |
| pMJ019 | Protein purification, activity assays *in vitro* and *in vivo* | Derivative of pLG012. Contains a Strep-tag II-encoding sequence fused downstream to the *drt4* gene (F4A mutant) | Chloramphenicol |

| | | | |
|---|---|---|---|
| pMJ020 | Protein purification, activity assays *in vitro* and *in vivo* | Derivative of pLG012. Contains a Strep-tag II-encoding sequence fused downstream to the *drt4* gene (N521A mutant) | Chloramphenicol |
| pMJ021 | Protein purification, activity assays *in vitro* and *in vivo* | Derivative of pLG012. Contains a Strep-tag II-encoding sequence fused downstream to the *drt4* gene (F523A mutant) | Chloramphenicol |
| pMJ022 | Protein purification, activity assays *in vitro* | pRSF-Duet1 (Novagen, Sigma-Aldrich 71341-3) derived plasmid encoding the *ncRNA-gp44* C-terminal Strep tag II. | Kanamycin |
| pMJ023 | Protein purification, activity assays *in vitro* | pRSF-Duet1 (Novagen, Sigma-Aldrich 71341-3) derived plasmid encoding the *ncRNA-gp44* (D152G mutant) C-terminal Strep tag II. | Kanamycin |
| pMJ024 | Activity assays *in vitro*, protein purification | pBAD plasmid encoding the *ncRNA-gp44* fused to C-terminal His tag II. | Ampicillin |
| pMJ025 | Activity assays *in vitro*, protein purification | pBAD plasmid encoding the *ncRNA-gp44* (D152G Bas17 escaper mutant) fused to C-terminal His tag II. | Ampicillin |

**Supplementary Tab. 2: Oligonucleotides**

| Name | Sequence (5'-3') | Purpose |
|---|---|---|
| MJ508 | CCCCCCCCCCCCCCCCCCCC | Oligonucleotide for ssDNA sequencing preparation |
| (U)5G(U)5 | [56-FAM]UUUUUGUUUUU[3IABkFQ] | Substrate for DRT4 RNase activity assay |

**Supplementary Tab. 3: Bacterial strains**

| Strain | Genotype | Purpose | Source |
|---|---|---|---|
| *E. coli* MG1655 ΔRM | *F– λ– ilvG– rfb-50 rph-1 Δmrr-hsdRMS-mcrBC ΔmcrA* | BASEL library screen | Maffei et al. [4] |
| *E. coli* MG1655 | *K-12* $F^-$ $\lambda^-$ $ilvG^-$ *rfb-50 rph-1* | *E. coli* growth assay | Department of Protein-DNA interactions, Life Science Centre, Vilnius University |
| *E. coli* DH5*α* | *F– Φ80lacZΔM15 Δ(lacZYA-argF) U169 recA1 endA1 hsdR17 (rK–, mK+) phoA supE44 λ– thi-1 gyrA96 relA1* | Cloning, expression | Thermo Scientific (EC0112) |
| *E. coli* BL21 Star (DE3) | *F- ompT hsdSB (rB- mB-) gal dcm rne131 (DE3)* | Expression | Invitrogen (C601003) |

**Supplementary Tab. 4: Cryo-EM data collection, refinement and validation statistics**

| | **DRT4** (EMD-59845, PDB-ID: 34GY) | **DRT4 complex with dApCpp** (EMD-59846, PDB-ID: 34HC) | **DRT4 complex with gp44** |
|---|---|---|---|
| *Data collection and processing* | | | |
| **Magnification** | ×92000 | | |
| **Voltage (kV)** | 200 | | |
| **Exposure (e⁻/Å$^2$)** | 30 | | |
| **Defocus range (µm)** | -2.2 to -1.0 | | |
| **Pixel size (Å)** | 1.1 | | |
| **Initial particles** | 1,708,489 | 2,041,746 | 1,623,810 |
| **Final particles** | 1,210,288 | 1,162,671 | 101,808 |
| **Symmetry imposed** | D3 | D3 | C1 |
| **Resolution (Å)** | 2.23 | 2.37 | 6.32 |
| **FSC cut-off** | 0.143 | 0.143 | 0.143 |
| **Map resolution range (Å)** | 2.2-4.8 | 2.3-5.0 | 6.2-20 |
| *Refinement* | | | |
| **Model resolution (Å)** | 2.9 | 2.7 | - |
| **0.143 FSC Threshold (Å)** | 2.2 | 2.2 | - |
| **Map sharpening *B* factor** | 110.7 | 135.8 | - |
| **Model composition** | | | |

| | | | |
|---|---|---|---|
| Non-hydrogen atoms | 25,998 | 26,778 | - |
| Protein residues | 3030 | 3102 | - |
| Nucleotides | 54 | 54 | - |
| Ligands | 0 | MG: 6<br>F2A: 6 | - |
| ***B* factors (Å$^2$)** | | | |
| Protein | 105.09 | 79.48 | - |
| Nucleotide | 157.11 | 121.25 | - |
| Ligands | - | 103.90 | - |
| *Validation* | | | |
| **Bonds (r.m.s. deviations)** | | | |
| Length (Å) (# > 4s) | 0.003 (0) | 0.004 (0) | - |
| Angles (°) (# > 4s) | 0.527 (11) | 0.642 (6) | - |
| **MolProbity score** | 0.82 | 0.95 | - |
| **Clash score** | 1.12 | 1.86 | - |
| **Poor rotamers (%)** | 0.00 | 0.00 | - |
| **Ramachandran plot (%)** | | | |
| Outliers | 0.00 | 0.00 | - |
| Allowed | 0.20 | 0.74 | - |
| Favored | 99.80 | 99.26 | - |
| **Rama-Z score, whole (r.m.s. Rama-Z)** | -0.60 (0.15) | -0.60 (0.14) | - |
| **CC (mask)** | 0.85 | 0.83 | - |

| | | | |
|---|---|---|---|
| **CC (box)** | 0.87 | 0.76 | - |

#### Supplementary References

1. Gao, L. *et al.* Diverse enzymatic activities mediate antiviral immunity in prokaryotes. *Science* **369**, 1077–1084 (2020).

2. Cernooka, E., Rumnieks, J., Tars, K. & Kazaks, A. Structural Basis for DNA Recognition of a Single-stranded DNA-binding Protein from Enterobacter Phage Enc34. *Sci Rep* **7**, 15529 (2017).

3. Rong, X. *et al.* DNA polymerization activates RNA cleavage of a reverse transcriptase-like antiviral enzyme. *Science* **393**, eaef3178 (2026).

4. Maffei, E. *et al.* Systematic exploration of Escherichia coli phage-host interactions with the BASEL phage collection. *PLoS Biol* **19**, e3001424 (2021).
